# The BTB-ZF transcription factor Tramtrack69 shapes neural cell lineages by coordinating cell proliferation and cell fate

**DOI:** 10.1101/338285

**Authors:** Françoise Simon, Anne Ramat, Sophie Louvet-Vallée, Jérôme Lacoste, Angélique Burg, Agnès Audibert, Michel Gho

**Affiliations:** CNRS, IBPS, UMR 7622, Laboratory of Developmental Biology, Paris F-75005, France.; Sorbonne Université, UMR 7622, Laboratory of Developmental Biology, Paris F-75005, France.

## Abstract

Cell diversity in multicellular organisms relies on coordination between cell proliferation and the acquisition of cell identity. The equilibrium between these two processes is essential to assure the correct number of determined cells at a given time at a given place. Here, we show that Tramtrack-69 (Ttk69, a BTB-ZF transcription factor ortholog of the human PLZF factor) plays an essential role in controlling this balance. In the *Drosophila* bristle cell lineage, producing the external sensory organs composed by a neuron and accessory cells, we show that *ttk69* loss of function leads to supplementary neural-type cells at the expense of accessory cells. Our data indicate that Ttk69 (1) promotes cell-cycle exit of newborn terminal cells by downregulating *cycE*, the principal cyclin involved in S-phase entry and (2) regulates cell fate acquisition and terminal differentiation by downregulating the expression of *hamlet* and upregulating that of *Suppressor of Hairless*, two transcription factors involved in neural-fate acquisition and accessory-cell differentiation, respectively. Thus, Ttk69 plays a central role in shaping neural cell lineages by integrating molecular mechanisms that regulate progenitor cell-cycle exit and cell-fate commitment.

**Summary statement:** Tramtrack-69, a transcription factor orthologous to the human tumor-suppressor PLZF, plays a central role in precursor cell lineages by integrating molecular mechanisms that regulate progenitor cell-cycle exit and cell-fate determination.

## Introduction

Organisms are composed of morphologically and functionally distinct cell types. Such cell diversity is generated from a restricted set of precursor cells producing a limited number of differentiated cells. Division of precursor cells gives rise to daughter cells that differentiate and acquire specific fates. The transit from a proliferative to cell-cycle arrested state during this process is tightly regulated and requires changes in transcriptional programs. Disentangling the molecular mechanisms that control the balance between proliferation and differentiation is essential for understanding the formation and maintenance of organisms, as well as human diseases, such as cancer, in which this process is disturbed.

BTB-ZF transcription factors are involved in a wide variety of biological processes (Kelly and Daniel, 2006). They include *Drosophila* Broad-complex factors (BR-C), Bric-à-brac (Bab), and several pox virus zinc-finger proteins (Chaharbakhshi and Jemc, 2016). All possess a protein/protein interaction motif (BTB/POZ) at the N-terminus that allows protein homo and multimerization and one or several zinc-finger DNA-binding motifs (Bonchuk et al., 2011). These proteins are conserved from *Saccharomyces cerevisiae* to *Homo sapiens* and act as transcriptional repressors or activators, depending on the BTB domain (Siggs and Beutler, 2012). The founding BTB-ZF members are all *Drosophila* transcriptional repressors that regulate processes such as metamorphosis, ovary development, and neurogenesis(Guo et al., 1995; Karim et al., 1993; Sahut-Barnola et al., 1995). In vertebrates, the human BTB-ZF, promyelocytic leukemia zinc finger (PLZF), acts as a tumor-suppressor maintaining cell growth inhibition and quiescence by transcriptional repression of the *c-myc* proto-oncogene (McConnell et al., 2003). Accordingly, *plzf* loss of function has been correlated with prostate and lung cancer (Jin et al., 2017). Moreover, this factor regulates organogenesis by controlling the balance between self-renewal and differentiation of neural stem cells (Gaber et al., 2013; Sobieszczuk et al., 2010). Overall, BTB-ZF proteins have fundamental and conserved roles during development, controlling cell proliferation and differentiation.

The *Drosophila* ortholog of PLZF, Tramtrack (Ttk), also plays multiple roles during development, including cell proliferation and cell-fate decisions in the nervous system, intestinal stem cells, photoreceptors, and tracheal cells (Araujo et al., 2007; Badenhorst, 2001; Giesen et al., 1997; Lai and Li, 1999; Liu et al., 2016; Wang et al., 2015). In particular, Ttk is a key regulator of cell fate in the peripheral nervous system, in which it promotes non-neural instead of neural fates (Guo et al., 1995). Ttk is considered to be a transcriptional repressor from the initial studies on *even skipped* and *fushi tarazu* genes(Brown et al., 1991; Harrison and Travers, 1988; Harrison and Travers, 1990). The *ttk* locus encodes two proteins, Ttk69 and Ttk88, via alternative splicing (Read and Manley, 1992). Both isoforms share a common conserved N-terminal BTB/POZ domain but contain divergent C-terminal zinc-finger C2H2-type domains for DNA binding, conferring specific DNA binding and probably independent functions for each isoform(Read and Manley, 1992). Both have specific functions during development, although Ttk69 appears to have a broader spectrum of functions than Ttk88. For example, during eye development, Ttk69, but not Ttk88, is expressed in all photoreceptor cells during the pupal stage and promotes specific non-neuronal fates, such as cone cells (Lai and Li, 1999). Similarly, Ttk69, but not Ttk88, is expressed in the embryonic nervous system, where it is required for proper glial cell development (Giesen et al., 1997). In the intestine stem cell lineage, *ttk69* loss of function leads to re-specification of enteroblasts into enteroendocrine cells, whereas *ttk88* loss of function has no phenotype (Wang et al., 2015). In addition to its well-known role in cell identity, Ttk has been also shown to be involved in cell-cycle regulation. More precisely, it has been shown that overexpression of Ttk69, but not Ttk88, causes the complete loss of mitosis in the eye disc morphogenetic furrow through the repression of the expression of String, the positive regulator of the G2/M transition (Baonza et al., 2002). Similarly, a significant increase of mitotic cells is observed in intestinal *ttk69* mutant clones, indicating that Ttk69 negatively regulates intestinal stem-cell proliferation (Wang et al., 2015). Altogether, these data highlight the essential role of Ttk69, but not Ttk88, on the control of cell proliferation and the acquisition of cell fate. In mechanosensory organs, the loss of both Ttk isoforms leads to complete transformation of sensory cells into neurons (Guo et al., 1995). Such extreme cell transformation prevents further studies to reveal elusive effects of Ttk isoform on cell fate determination and other biological processes, such as cell differentiation and cell proliferation. Indeed, our previous studies have shown that the loss of Ttk69 alone also induces cell proliferation (Audibert et al., 2005). This suggests that the study of mutations affecting specificaly Ttk69 may reveal its cryptic roles that are not observed when analyzing complete *ttk* loss of function mutations. Here, we focus on how Ttk69 controls the balance between cell proliferation and the acquisition of cell fate in the bristle system.

The *Drosophila* external mechanosensory organs, or bristles, are an excellent model system to study the balance between proliferative and determined states of progenitor cells (Fichelson et al., 2005). Each bristle is composed of a shaft and an annular cuticular structure, called the socket, at its base. At the cellular level, only four specialized cells, with a common origin, compose this relatively simple structure: two outer cells, the socket and shaft cells, and two inner cells, the neuron and sheath cell (Hartenstein and Posakony, 1989). Each cell differs from the other by its size, relative position, and expression of specific markers (Figure 1A). They arise from the division of a primary precursor cell (or pI) after a stereotypical sequence of four asymmetric cell divisions (the bristle cell lineage). In the dorsal thorax, pI cells divide to generate a posterior secondary precursor cell (pIIa) and an anterior secondary precursor cell (pIIb). The division of pIIa leads to the formation of the outer cells (the pIIa sub-lineage), whereas the pIIb cell gives rise to the inner cells (the pIIb sub-lineage), following two rounds of division. First, pIIb divides to give rise to a glial cell that enters apoptosis shortly after birth and a tertiary precursor cell, pIIIb. Then, pIIIb divides to produce the sheath and the neuron (Fichelson and Gho, 2003; Gho et al., 1999). At each of these divisions, the Notch (N) pathway is differentially activated in only one daughter cell. This differential activation ensures the acquisition of different fates by both daughter cells (Guo et al., 1996). As such, the N-pathway does not specify particular identities, but its activation triggers different outcomes depending on the cellular context, likely in cooperation with other factors that specify cell fate (Ramat et al., 2016). Only some of these specific factors are known in the bristle lineage. Two are Sequoia (Seq) and Hamlet (Ham), two zinc-finger transcription factors, expressed in pIIb sub-lineage cells, that have a critical role in the acquisition of inner cell identity (Andrews et al., 2009; Moore et al., 2002; Moore et al., 2004). Indeed, sensory organs (SO) in *ham* and *seq* mutants are composed of external cells only, due to respecification of the inner cells. Moreover, it has also been shown that Seq controls *ham* expression, indicating that these factors are related in a complex regulatory network of transcription factors (Andrews et al., 2009).

**Figure 1.**
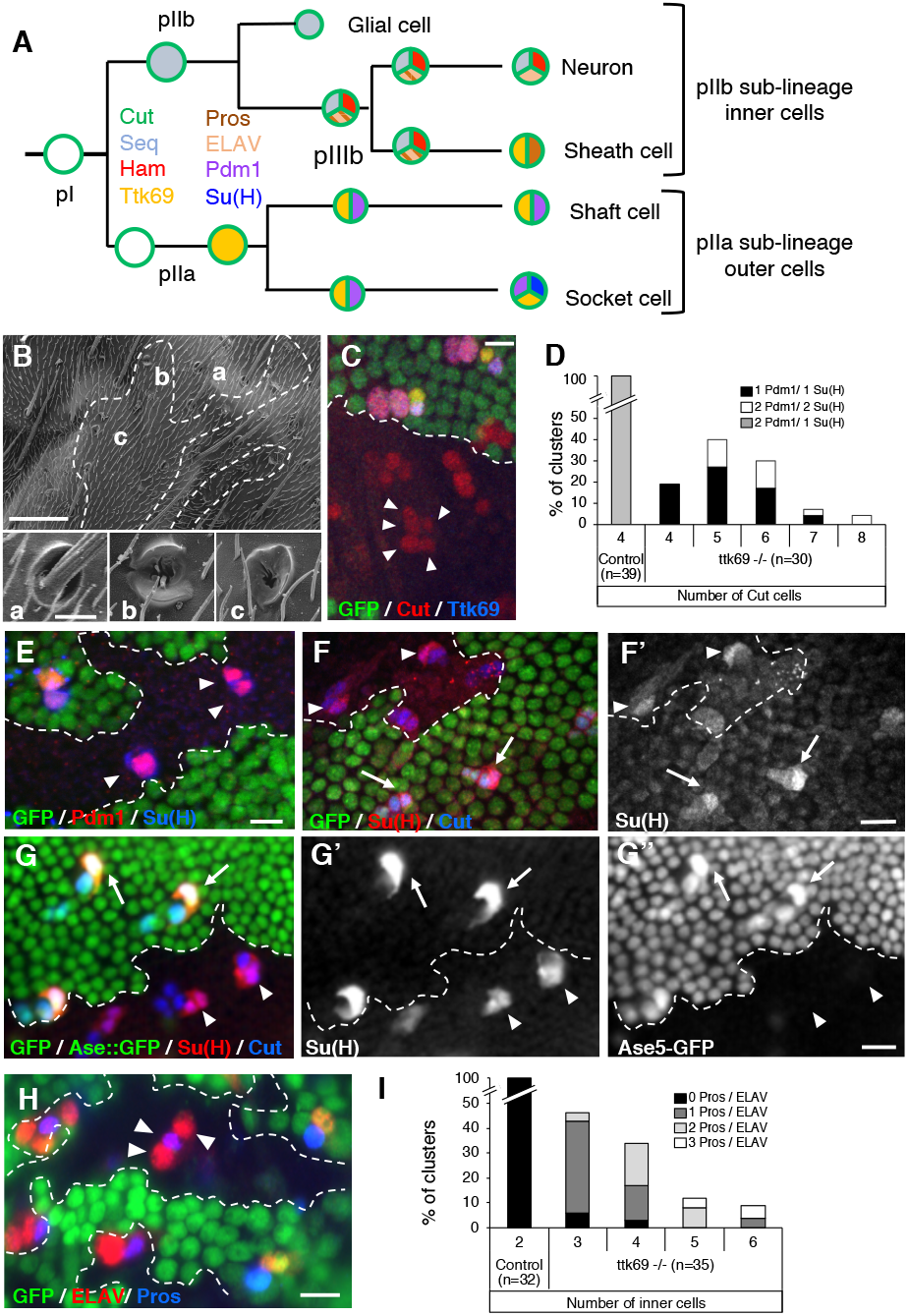
Ttk69 loss of function leads to extra-terminal cells in SO. (A) Scheme of the wild type bristle lineage. Cells are represented by circles and cell markers by specific colors. (B) Scanning electron micrograph of Ttk69 SO at the external level. The Ttk69 clone is outlined by a white dashed line. Control (a) and Ttk69-mutant SO (b and c). (C, E-H) Ttk69-mutant SO at the cellular level. Ttk69 clones were detected by the absence of GFP and outlined by a white dashed line. Pupae were at 28 h APF except in F where they were at 23 h APF. (C) Ttk69-mutant SO were composed of more than four cells (arrowheads). Sensory cells were identified by Cut (red) and Ttk69 (blue) immunoreactivity. (D) Histogram showing the percentage of SO harboring four to eight cut positive cells in control and in mutant SO. The percentage of SO with different combinations of Pdm1- and Su(H)-positive cells is shown (Black bar, only one Pdm1/Su(H)-positive cell. White bar, two Pdm1/Su(H) positive cells. Gray bar, 1 Su(H)-positive cell among 2 Pdm1 positive cells). (E) Outer cells acquired a socket fate: All outer cells (specifically marked by Pdm1 (red)) expressed the socket marker Su(H) (blue) in Ttk69-mutant SO (arrowheads). (F-G) Auto-amplification of Su(H) was impaired in Ttk69-mutant SO. (F, F’) In 23 h APF pupae, accumulation of Su(H) protein (F’ and red in F) was similar in Ttk69-mutant SO (arrowheads) and in control SO (arrows). (G-G”) In 28 h APF old pupae, Su(H) immunoreactivity (G’ and red in G) and Su(H) autoamplification assessed by the ASE5::GFP reporter (G’’ and green in G). Note that Su(H) auto-amplification was absent. (H) More than two inner cells (arrowheads) are present in Ttk69-mutant SO. Inner cells are revealed by ELAV (red) and Pros (blue) immunoreactivity. (I) Histogram showing the percentage of SO harboring two to six pIIb cells in control and in mutant SO. The percentage of clusters with zero (black bars), one (dark grey bars), two (light grey bars), or three (white bars) inner cells positive for both Pros and ELAV immunoreactivity (Pros/ELAV) is indicated. Scale bars: 50 μm in B, inset 5 μm; 10 μm in C, E, F, G, H.

Here, we use the bristle lineage to explore how Ttk69 coordinates terminal cell determination and cell-cycle arrest. We show that loss of *ttk69* leads to the production of supernumerary progenitor cells and the re-specification of cell fate identity. Notably, we observed a cell transformation in which an outer cell acquired an inner cell precursor identity, precisely the pIIa shaft cell adopts a pIIIb cell fate. Since outer and inner cells are cousin cells, this kind of transformation has been previously named cousin-cousin cell transformation (Moore et al., 2004). We identified the *cycE* gene, encoding the essential cyclin required for entry into S-phase, as a Ttk69 downstream gene. In addition, we show that Ttk69 regulates cell-fate acquisition and terminal differentiation by controlling the expression of *ham* and *Suppressor of Hairless (Su(H))*, which encodes the transducing transcription factor of N-receptor signaling. We propose that Ttk69 is a central node of a transcriptional regulatory network that assures cell lineage completion by controlling the acquisition of terminal cell fates and the arrest of cell proliferation.

## Results

### Ttk69 loss of function leads to the formation of SOs with extra inner cells and only one type of outer cells

To precisely determine the involvement of Ttk69 in cell-cycle progression and cell-fate determination, we studied somatic clones of *ttk*^1e11^, which specifically disrupts Ttk69, hereafter called Ttk69 clones (Lai and Li, 1999). SOs inside Ttk69 clones (called Ttk69-mutant SOs) were devoid of the shaft and presented only sockets externally (Figure 1B). At the cellular level, 82% (n = 30) of the mutant organs were composed of more than four cells (up to eight cells) at 28 h after pupal formation (APF) (Figures 1C arrowheads and D). Among the SO cells, one (67%) or two (33%) cells were Pdm1-positive outer cells (n = 30) and, in all cases, they expressed Su(H), a landmark of socket cells (Figures 1D and E). These data show that the absence of the shaft structure is associated with the lack of a cell expressing a shaft signature (Pdm1 positive, Su(H) negative). We observed that in Ttk69-mutant SO socket cells, although present, did not have a normal shape (Figure 1B, insets b, c). It was previously shown that the normal socket cell differentiation is dependent of the Notch transcription factor Su(H) auto-amplication. Thus, Su(H) is first induced in presumptive socket cells in response to the N-pathway and subsequently boosted via its binding to a 3’-enhancer (ASE5, (Barolo et al., 2000; Liu and Posakony, 2014)). Initially, at 23 h APF, Su(H) expression was similar in control and in Ttk69-mutant socket cells (Figures 1F, F’, quantified in Figure S1A). Later, at 28 h APF, using a ASE5::GFP reporter to assess Su(H) autoamplification, we failed to detect a GFP signal in Ttk69-mutant socket cells indicating that Su(H) amplification had not occurred (Figures 1G-G”, compare GFP expression in control socket cells, arrows, with Ttk69-mutant socket cells, arrowhead). This was associated with no further increased in Su(H) level in Ttk69-mutant socket cells (Figure 1G’, compare Su(H) accumulation in control socket cells, arrows, with Ttk69-mutant socket cells, arrowhead, quantified in Figure S1A). Since the initial Su(H) expression was not affected in the absence of Ttk69, this suggests that this factor is not involved in socket cell determination. However, since Su(H) auto-amplification was impaired in a Ttk69-mutant background, our data indicate that it is indeed involved in socket cell terminal differentiation.

The remaining cells in Ttk69-mutant SOs expressed inner-cell markers. Immunostaining against ELAV and Prospero (Pros) revealed the presence of one to four neurons and one or no sheath cell (Figure 1H). We also observed up to three cells per cluster (n = 35) that were positive for both ELAV and Pros immunostaining (Figures 1H and I). This suggests the presence of either additional pIIIb precursor cells or post-mitotic cells in which the fate was not yet well resolved or not at all (Ramat et al., 2016). Thus, the Ttk69-mutant lineage is probably not yet completed at 28h APF. Overall, these data show that loss of function of Ttk69 leads to the formation of SOs composed of extra inner cells and only socket cells as the outer cell type.

### Ttk69 promotes cell-cycle arrest and triggers terminal cell-fate identities

Several non-exclusive explanations can account for the presence of SOs with supplementary cells in Ttk69-mutant SOs. One is that the glial cells do not die but divide and produce extra terminal cells. We examined this possibility by studying cell death in Ttk69-mutant SOs. TUNEL assays showed that glial cells undergo apoptosis in Ttk69-mutant SOs at the same time as in control organs located outside the mutant clone (Figure 2A, n = 3). Thus, the supplementary cells do not originate from glial cells that resume proliferation. It is also possible that supplementary cells arise from bristle cells that do not properly exit from the cell cycle and continue to proliferate. We explored this possibility by searching for metaphasic cells using phospho-ser10 histone-3 (PH3) immunoreactivity at 28 h APF, when control SO cells are already post-mitotic. Ttk69-mutant SOs containing four or more cells harbored sensory cells positive for PH3 (Figure 2B, arrowheads, 20% of SOs, n = 25). As these clusters contained the terminal number of cells, these data show that the metaphasic cells were not due to delayed divisions. Thus, these data indicate that Ttk69-mutant SOs harbor supplementary cells due to additional mitoses.

**Figure 2.**
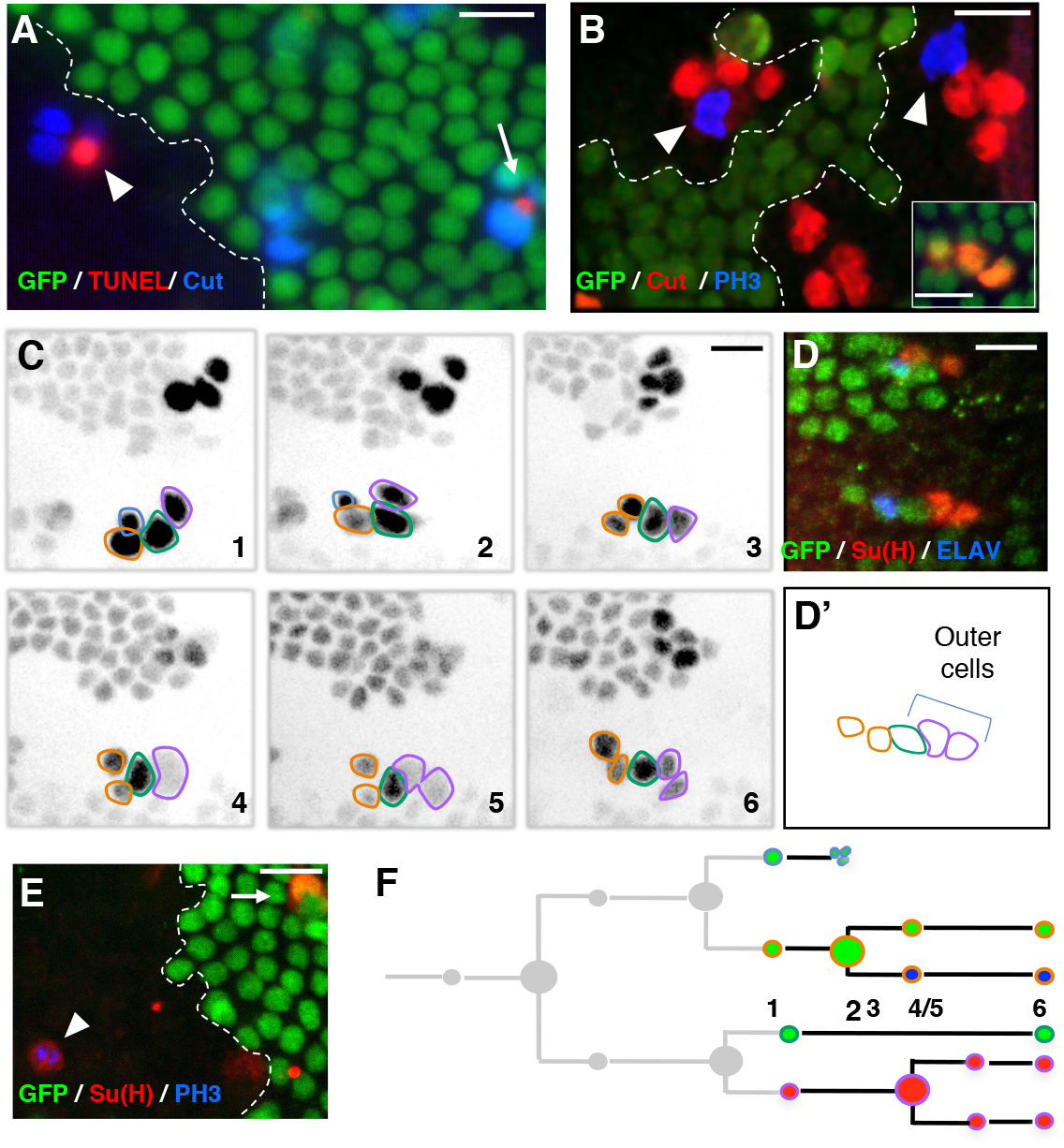
Extra mitoses in Ttk69-mutant socket cells. (A, B, E) Ttk69 clones, outlined by a white line are detected by the absence of GFP (green) in fixed tissues. (A) Apoptosis occurred at the same time in mutant (arrowhead) and control (arrow) SO. Sensory cells (blue); apoptotic cells (TUNEL staining, red). (B) Extra mitoses (PH3 immunoreactivity (blue), arrowheads) in Ttk69-mutant SO composed of four cells (red); control cluster in the inset. (C, D) Correlative 4D live imaging and lineage analysis showing an extra division of socket cells. A Ttk69 clone was identified by the lack of GFP expression in epithelial cells and SO inside the clones were imaged. (C) Representative frames (1-6), depicted in inverted fluorescence, from a time-lapse recording of one Ttk69-mutant SO at 19 h APF. Glial cell outlined in blue, pIIb and its progeny in orange, the shaft cell in green, and the socket cell and its daughter in purple. Frames 4 and 5, division of the socket cell. Apoptosis of the glial cell between frames 2 and 3. (D, D’) Immunostaining and schematic representation of the same cluster after the time-lapse recording shown in C. Non-clonal epithelial and sensory cells (GFP expression, green), neurons (ELAV, blue) and socket cells (Su(H), red) immunoreactivity. (E) Cell divisions in full-determined socket cells. PH3 (blue) and Su(H) (red) immunoreactivity was detected in the same cell (arrowhead); Control socket cell (arrow). (F) Schematic view of the lineage shown in C. Cells are encircled using the same color code as in C and filled with the same color as in D. Scale bars: 10 μm.

To unambiguously define the origin of the supernumerary cells, we used correlative microscopy that combines live imaging to record the entire pattern of cell divisions in Ttk69-mutant SOs followed by immunolabeling to identify cell identities. During time-lapse recording, sensory cells were identified by the expression of GFP under the control of the *neuralized* promoter (neur-GFP). At the end of each recording, the imaged notum was fixed and immunolabeled with anti-Su(H), as well as anti-ELAV and anti-Pros to highlight outer and inner cells, respectively. Imaged SOs could be unambiguously recognized within the fixed nota by their relative position with respect to the midline, the position of the macrochaetae, or the rows of microchaetae (Fichelson and Gho, 2004). We confirmed the presence of extra cell divisions and revealed an unexpected cell transformation event. First, there was a supplementary division in a pIIa daughter cell, identified as the future socket cell by its position in the cluster (Figure 2C, panels 4 and 5). This extra division was symmetric, leading to two Su(H)-positive socket cells (Figure 2D, D’). We also observed socket cells in mitosis, identified by Su(H) and PH3 immunoreactivity, in fixed material (Figure 2E, arrowhead, n = 3). Thus, future socket cells undergo an extra division in the absence of Ttk69 (Figure 2F). In addition, the anteriorly located pIIa daughter cell, the presumptive shaft cell that normally does not divide, underwent repetitive cell divisions (Figure 3A, panels 3 and 5). Surprisingly, immunostaining of the resulting clusters showed that cells arising from these extra divisions acquired a neural fate, as they expressed Pros and ELAV (Figure 3B, B’), two markers expressed in sheath and neuron cells, respectively. This suggests that presumptive shaft cells underwent cousin-cousin cell transformation in which outer cells acquired an inner cell fate (Figure 3C). Consistent with this possibility, we also observed cells with weak expression of Pdm1 associated with weak expression of Pros in fixed material, suggesting that they were midway through transformation (arrowhead in Figures 3D, D’, D”). Furthermore, we detected clusters harboring two Pros-positive cells, of which one was dividing (PH3 positive, arrowhead in Figures 3E, E’, E”). These different lines of evidence led us to conclude that the presumptive shaft cells underwent cell fate re-specification and acquired a pIIIb precursor cell identity. We never observed by *in vivo* recordings cell lineages in which both pIIa daughter cells entered division. This is likely due to the low probability of such cases. We do not favor the possibility that the division of pIIa daughter cells is mutually exclusive, as using fixed material, we observed SOs composed of more than five cells and harboring two socket cells, a situation that required ectopic division of both pIIa daughter cells.

**Figure 3.**
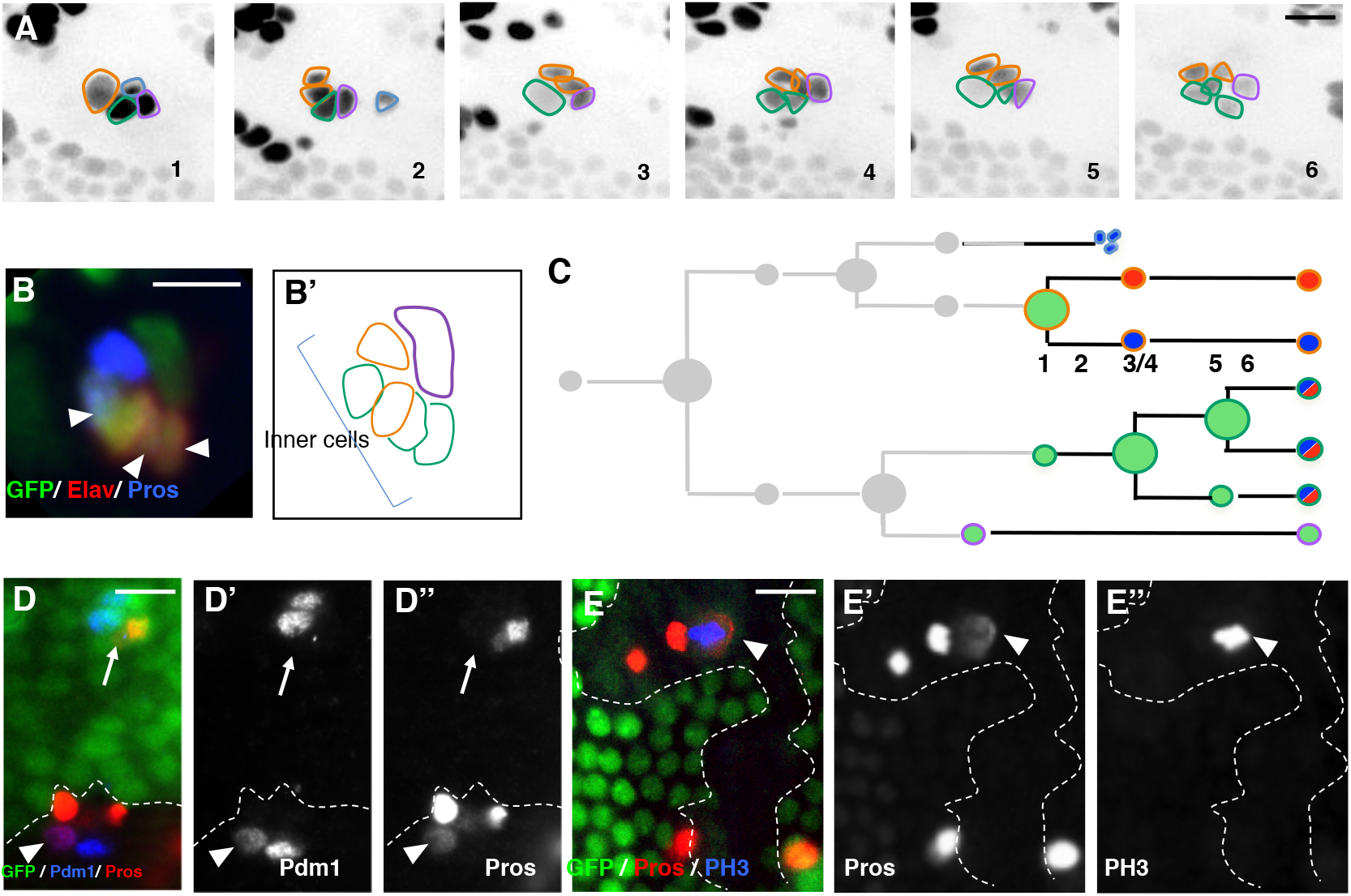
Presumptive Ttk69-mutant shaft cells undergo cell transformation toward inner precursor cells. (A) Correlative 4D live imaging and lineage analysis showing extra divisions of the presumptive shaft cell. Representative frames (1-6) depicted in inverse fluorescence from a time-lapse recording of one Ttk69-mutant SO at 20 h APF. Cells outlined as in Figure 2C. Frame 2: apoptosis of the glial cell. Frames 3 and 5: two rounds of division of the presumptive shaft cell. (B, B’) Immunostaining and schematic representation of the same cluster after the time-lapse recording. Sensory cells in green. Inner-cells identified by ELAV (red) and Pros (blue) immunoreactivity. (C) Schematic view of the lineage shown in A. Cells are encircled using the same color code as in A and filled with same colors as in B. (D, E) Ttk69-mutant clones detected by the absence of GFP (green) are outlined by a white line. (D-D”) Cousin-cousin cell fate transformation. Arrowhead, mutant cell having both Pdm1 (D’ and blue in D) and Pros (D” and red in D) immunoreactivities (outer and inner cell marker respectively), situation never observed in control (arrow). (E-E”) Extra mitoses of inner cells revealed by PH3 immunoreactivity (E” and blue in E). Inner-cells identified by Pros immunoreactivity (E’ and red in E). Scale bars: 10 μm.

The re-specification of presumptive shaft cells into inner precursor cells can explain, in part, the observation that Ttk69-mutant sensory clusters harbored up to six neural (ELAV/Pros positive) cells, showing that extra inner-cells originate at the expense of outer cells. Overall, these results suggest that Ttk69 promotes cell precursor exit, acting both by arresting cell proliferation and triggering terminal cell fate identity.

### Ttk69 induces cell cycle exit via transcriptional repression of *cycE* expression

Our results suggest that Ttk69 regulates cell proliferation. We have already shown that Ttk69-mutant sensory cells strongly accumulate CycE protein (Audibert et al., 2005). We studied whether supplementary mitosis related to the accumulation of ectopic CycE by assessing whether a reduction in the dose of CycE could revert the phenotype of supplementary cell divisions observed in Ttk69-mutant SOs. As already described, we observed that most SOs inside Ttk69-mutant clones contained more than the normal four cells (85%, n = 42, Figure 4A). This dropped to 18% when the clones were induced in a *cycE^AR95/+^* heterozygous background (n = 35). In another set of experiments to study the number of socket cells, 45% of Ttk69-mutant SOs harbored duplicated Su(H)-positive socket cells (n = 55), whereas only 11% did so in the *cycE^AR95/+^* heterozygous background (n = 51). These results show that reducing the dose of CycE is sufficient to markedly reduce the number of supplementary divisions observed in Ttk69-mutant SOs. This strongly suggests that the supplementary mitoses observed in Ttk69-mutant SOs are mainly driven by the increase in CycE levels induced after Ttk69 loss of function.

**Figure 4.**
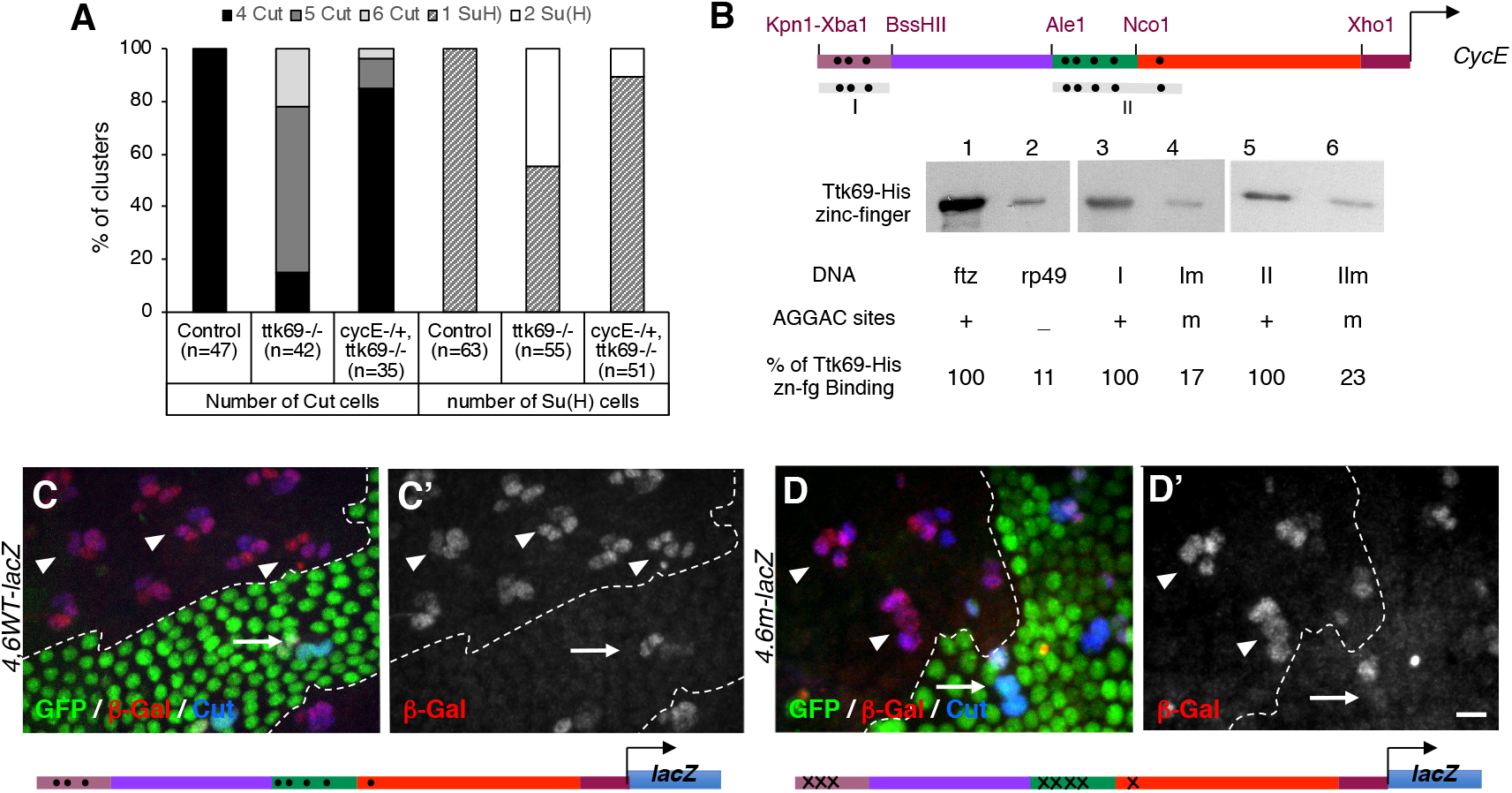
*cycE* expression is transcriptionally repressed by Ttk69. (A) The integrity of Ttk-mutant SO was rescued under *cycE heterozygous conditions.* Histogram showing the percentage of SO harboring four (black bars) or more (gray bars) Cut-positive cells (left) and one (hatched bars) or two (white bars) Su(H)-positive cells (right) located outside (control) and inside Ttk69 clones in *cycE^+/+^* or *cycE^AR95/+^* heterozygous backgrounds. (B) DNA-mediated Ttk69-His-zinc-finger pull-down assay. (Top) Diagram of Ttk69 binding sites (black dots) in *cycE* promoter. (Bottom) Magnetic beads were coated with: lane 1, *ftz* promoter bearing AGGAC binding sites (positive control); lane 2, *rp49* a AGGAC-free promoter (negative control); lanes 3 and 5, I and II regions of the *cycE* promoter respectively; lanes 4 and 6, Im and IIm regions of the *cycE* promoter respectively, in which AGGAC binding sites were replaced by an unrelated TCGAC sequence. (C, D). Expression pattern of the *4,6WT-lacZ* (C, C’), *4,6m-lacZ* (D, D’) *cycE* transcriptional reporters in control SO (arrows) and Ttk69-mutant SO (arrowheads). Ttk69 clones outlined by a white line were detected by the absence of GFP (green). Sensory cells (Cut immunoreactivity, blue) and expression of *cycE* transcriptional reporters (β-Gal immunoreactivity, red). Scale bars: 10 μm for C, D.

We analyzed the role of Ttk69 in the transcriptional expression of *cycE* by first testing the capacity of Ttk69 to bind the *cycE* promoter. It has been previously shown that a 4.6 Kb proximal fragment of the whole *cycE* promoter is able to recapitulate *cycE* expression in the embryonic peripheral nervous system (Jones et al., 2000). We have identified eight AGGAC canonical Ttk binding sites in this 4.6 Kb fragment organized in two clusters: three in region I and five in region II (Figure 4B)(Harrison and Travers, 1988; Harrison and Travers, 1990). The binding of Ttk69 to these regions was assessed by a DNA-mediated Ttk pull-down assay using the Ttk69 C-terminal domain, containing the C2H2-type zinc-finger. The Ttk69 zinc-finger domain behaved as expected, since it was efficiently retained on beads coated with DNA fragments bearing canonical Ttk-binding sites *ftz*, as well as on regions I or II (Figure 4B, lanes 1, 3, and 5). In contrast, we observed low-level nonspecific retention when beads were coated with DNA free of Ttk binding sites *(rp49*, Figure 4B, compare lanes 1 and 2); quantification showed that binding to the *rp49* probe was reduced to 11% (n = 5) of that observed using the *ftz* probe. We also observed low-level nonspecific retention when the beads were coated with regions I or II in which the Ttk binding sites were mutated (Im and IIm respectively, Figure 4B, compare lane 3 with 4 and lane 5 with 6). Binding to the Im probe was reduced to 17% (n = 5) of that containing region I and binding to the IIm probe was reduced to 23% (n = 5) of that containing region II. As such, this *in vitro* approach suggests that the control of *cycE* expression by Ttk69 relies on the direct binding of this factor to canonical Ttk binding sites to the *cycE* promoter. We then test this possibility *in vivo* by lac-Z transcriptional reporter strategy. We followed in sensory cells *lacZ* expression under the control of a wild-type 4.6 *cycE* promoter fragment (4.6WT) and a 4.6 cycE fragment in which all 8 Ttk69 binding sites were mutated (4.6m in which the AGGAC sequence was replaced by the ACTGC sequence) (Figure 4). As expected, the 4.6WT fragment recapitulates CycE expression in adult bristle sensory cells (Audibert et al., 2005). Indeed, we observed β-Gal accumulation in inner cells, whereas it was only barely detected in pIIa daughter cells (Figures 4C, C’ arrow and Figure S2A, B). Furthermore, in Ttk69-mutant SOs, the expression of the 4.6WT construct was upregulated. We observed a particularly high level of β-Gal accumulation in both pIIa daughter cells (Figures 4C, C’ arrowheads. Quantifications in Figure S1B). Surprisingly, the expression pattern of the 4.6m transcriptional reporter, where the eight Ttk69 binding sites has been mutated, was not modify in wild-type SOs (Figures 4D, D’ arrow and S2C, C’). Moreover, expression of the 4.6m-lacZ construct was still upregulated in Ttk69-mutant SOs (Figures 4D, D’, arrowheads. Quantifications in Figure S1B). As such, although *in vitro* experiments show that Ttk69 binds *cycE* promoter, the *in vivo* results indicate that repression of *cycE* expression in sensory cells is not mediated through binding to the canonical AGGAC Ttk-binding sites present in the 4.6 fragment.

To accurately define which part of the *cycE* promoter is required for Ttk69 regulation, we divided the 4.6Kb *cycE* cis-regulatory fragment into four regions (A to D) and monitored the regulatory activity of constructs with these regions deleted in bristle sensory cells (Figures 5A and S2A). All constructs were inserted at the same locus (using ΦC31 integrase-based tools, except line D-lacZ) to avoid expression variations due to genomic environment. A *cycE* promoter construct bearing a deletion of region A and C (ΔAC-lacZ) as well as a deletion of region B (ΔB-lacZ) showed an expression pattern similar to that of the 4.6Kb-lacZ construct, indicating that the deletion of these regions did not remove the Ttk69 regulatory domain (Figures S2D, D’, E, E’). In contrast, deletion of region D (ΔD-lacZ) led to high levels of βGal accumulation in all sensory cells under control conditions (Figures 5B, B’, arrows and S2F, F’). Moreover, the expression pattern was similar in Ttk69-mutant SOs and control SOs outside of the clones (Figures 5B, B’, arrowheads. Quantifications in Figure S1C). These data show that the cis-regulatory sequence required for Ttk69-mediated down-regulation of *cycE* expression is located in the D-sequence. Next, we analyzed *lacZ* expression under the control of the fragment D (D-lacZ) in wild-type and in Ttk69-mutant SOs. Although a low level of β-Gal accumulation was observed with the D-lacZ construct, this construct behaves similarly as the 4.6WT-lacZ construct. β-Gal was barely detected in pIIa control cells (Figures 5C, C’ arrow and S2G, G’) while a high level of β-Gal accumulation was observed in mutant SOs (Figures 5C, C’ arrowhead. Quantification Figure S1C). These data confirm that this region is sufficient to drive Ttk69-mediated *cycE* regulation. Although, the D-fragment harbors a Ttk69 binding site at the more distal location, we do not believe that this binding site plays a role on the Ttk69-mediated control of *cycE* expression since Ttk69 continues to regulate the expression of reporter when this Ttk69 binding site was mutated (see above, Figures 4D, D’). A such, these data suggest that Ttk69 may control *cycE* expression by binding directly to unknown binding sites located in the D-fragment or indirectly either via another factor that recognizes the D-fragment or via the regulation of the expression of *cycE* transcriptional regulators.

**Figure 5.**
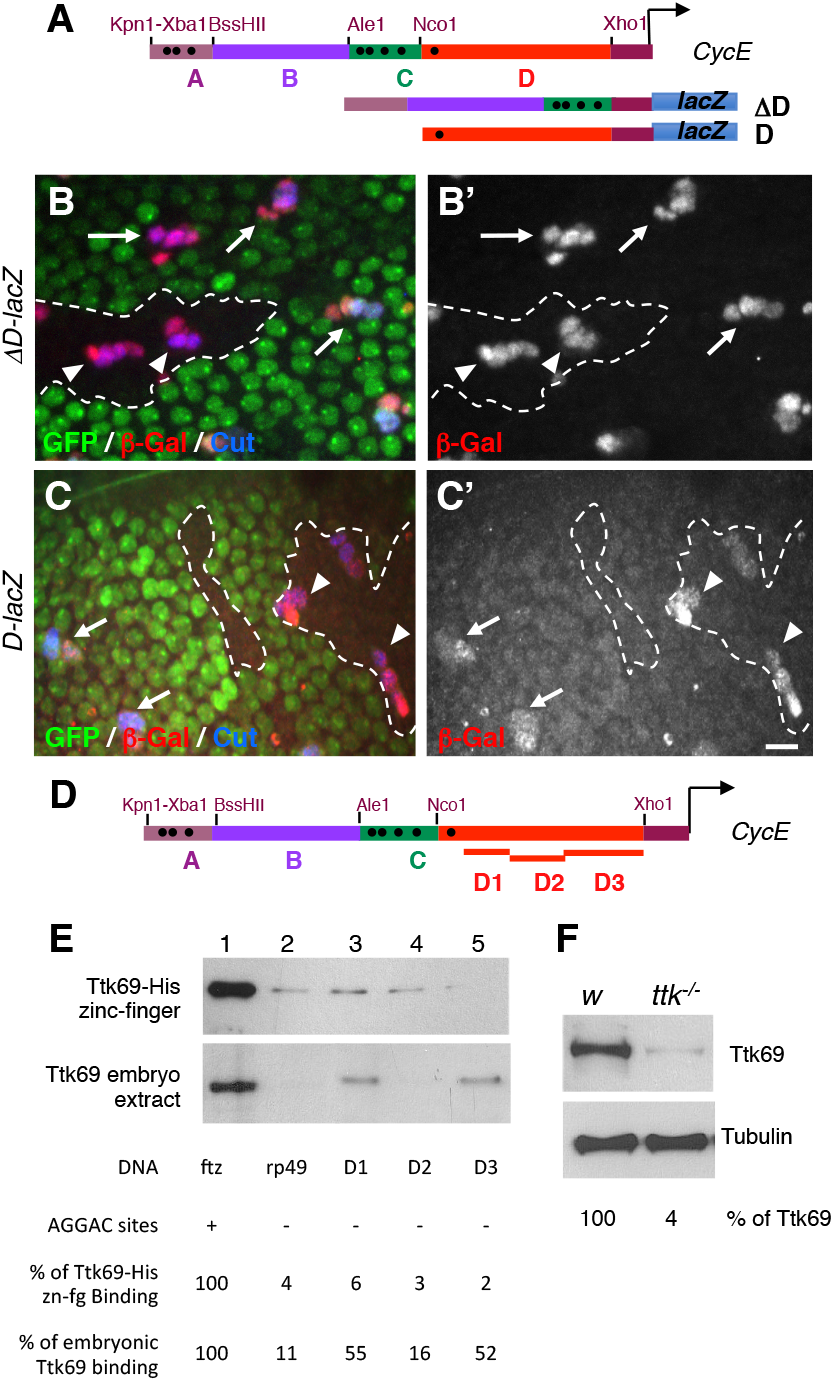
Ttk69 binds indirectly to the *cycE* promoter. (A) Diagram of *cycE* promoter depicting the four specific regions analyzed (A to D). Black dots, canonical AGGAC Ttk-binding sites. ΔD *cycE* and D *cycE* transcriptional reporters bearing a deletion of the D region and only the D region respectively. (B, C) Expression pattern of the *ΔD-lacZ* (B, B’) and *D-lacZ* (C, C’) in control SO (arrows) and Ttk69-mutant SO (arrowheads). Ttk69 clones outlined by a white line were detected by the absence of GFP (green). Sensory cells (Cut immunoreactivity, blue) and expression of *cycE* transcriptional reporters (β-Gal immunoreactivity, red and bottom panels). (D) As in (A) showing the regions of the fragment D analyzed (D1 to D3). (E) DNA-mediated pull-down assay using Ttk69-his-zinc-finger (top) and whole Ttk69 protein from an embryo protein extract (bottom). Magnetic beads were coated with: lanes 1 and 2, as in B; and lanes 3, 4, and 5, the D1, D2, and D3 regions, respectively, of the *cycE* promoter. (F) Detection of Ttk69 protein from a protein extract of one control *(white)* and one Ttk69-mutant embryo. Tubulin, loading control. Scale bars: 10 μm for B and C.

### Ttk69 binds to the *cjc,E*-promoter independently of canonical sites

To address these possibilities, we divided the D-fragment devoid of the Ttk69 binding site in three (D1-D3) and assessed their role in Ttk69 binding (Figure 5D). As expected, none retained the Ttk69 zinc-finger domain (Figure 5E, top panel, compare lanes 3 to 5 to the positive *fiz*-probe (lane 1) and the negative rp49-probe (line 2, n = 2). Next, we tested whether endogenous Ttk69 binds cycE promoter in association with other partners, by performing DNA-mediated Ttk pull-down assays using embryonic extracts and a specific Ttk69 antibodies (Figures 5E and F). Remarkably, although endogenous Ttk69 was not retained by the D2 cycE-promoter fragment (Figure 5E bottom panel, lane 4, quantification showed that binding to the *D2* probe was reduced to 16% (n = 5) of that observed using the *ftz* probe, which is similar to the 11% observed when we used the *rp49* probe (n = 5)), it was indeed retained on beads coated with the D1 and D3 fragments (Figure 5E, bottom panel, lanes 3 and 5, binding to the D1 probe was 55% and to the D3 probe was 52 % of that observed for the *ftz* probe (n = 5)) These data suggest that the binding of Ttk69 to D1 and D3 *cycE*-promoter fragments required either the native form of Ttk69 or other proteins present in embryonic extract.

### Ttk69 downregulates *hamlet* expression

We wished to identify Ttk target genes involved in cell fate regulation in SOs. We focused on two candidates, *hamlet (ham)* and *sequoia (seq)* (Andrews et al., 2009; Moore et al., 2004), because the phenotype associated with *ham* and *seq* loss of function, inner to outer cell transformation, is similar to that of *ttk* gain of function. In addition, *ham* and *seq* genes are expressed in patterns that are complementary to the expression pattern of *ttk* during the bristle lineage (Figures 1A and S3, see supplementary Figure 4 in (Andrews et al., 2009)). We thus studied the potential epistatic interactions between these three factors.

We first studied *ttk* expression in SOs when *ham* and *seq* were either overexpressed or downregulated. We used a temperature conditional driver to overexpress *ham* or *seq* late in development to avoid potential interference with outer-to-inner cell transformations induced by *seq* or ham-overexpression. Under these conditions, we detected no cell fate transformations, assessed by Su(H) immunoreactivity as a marker of outer socket fate (Figures S4A-C, Su(H) panels). This analysis revealed that Ttk69 expression was unaffected when either *ham* or *seq* were overexpressed (Figures S4A-C, Ttk panels. Quantification in Figure S1D). Reciprocally, we failed to observe modifications in the number of Ttk-positive cells early during development in *ham* or *seq* mutant SOs (20 and 22 h APF, Figures S4D-G). We observed sensory clusters harboring four Ttk69-positive cells only late in the bristle lineage in *seq* or *ham* mutant clones (24 h APF in 12% and 34% of clusters in *ham* and *seq* mutant clones, respectively, Figures S4D-E, F and G). These data suggest that supernumerary Ttk cells in *ham* and *seq* loss of function are due to cell transformation induced by the loss of function of *seq* or *ham*, rather than direct deregulation of *ttk69* expression. Thus, we conclude that *ttk* expression is independent of Ham and Seq.

Next, we analyzed whether *ham* or *seq* expression are controlled by Ttk69. We thus overexpressed Ttk69 in neurons, where it is never detected, and analyzed Seq and Ham protein accumulation. We used a similar strategy as before to overexpress Ttk69 late in development and observed no cell-fate transformation as shown by ELAV immunoreactivity (Figures 6A and B, ELAV panels). Under these conditions, Seq accumulated at the same level as in the control situation, whereas Ham immunoreactivity was strongly reduced (compare the right panels in Figures 6A and B for Seq and Ham detection; respectively. Quantification in Figure S1E, F). The observed effects were not due to changes in cell fate, as ELAV expression was unaltered. As such, we conclude that Ttk69 does not affect *seq* expression, whereas it downregulates *ham* expression.

**Figure 6.**
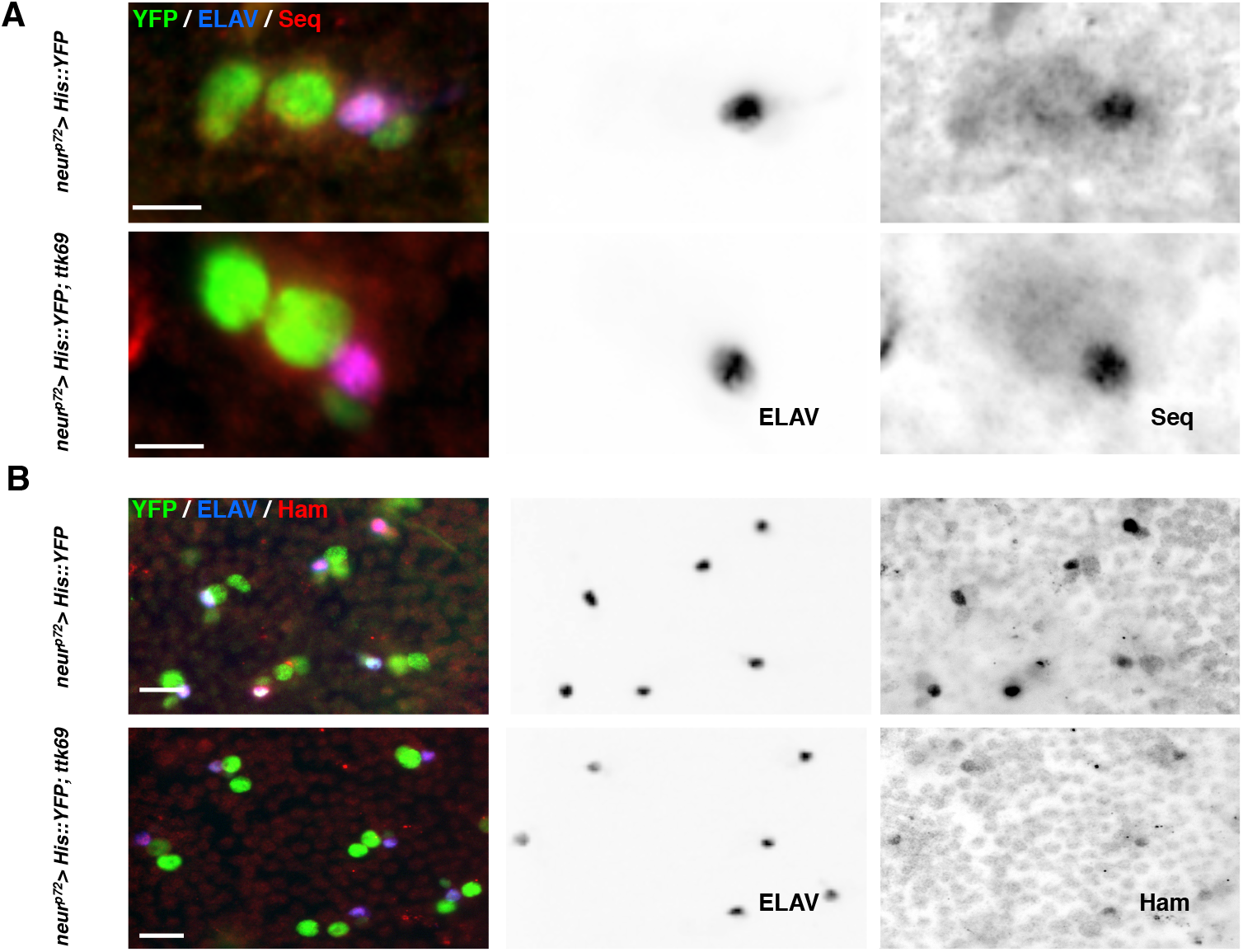
Ttk69 downregulates *hamlet* but not *sequoia* expression. (A, B) Ttk69 overexpression represses *hamlet* but not *sequoia* expression. Analysis of Seq (A) and Ham (B) protein accumulation after specific expression of Ttk69 in sensory cells. Sensory cells (green); neurons (blue), Ham and Seq (red). ELAV (middle panels), Ham, and Seq channels (right panels) are shown in inverted color. Scale bars: 5 μm in A and 10 μm in B.

### Ttk69 maintained non-neural cell fate via repression of *hamlet* expression

Our results show that Ttk69 downregulates *ham* expression. This suggests that *ham* is repressed in Ttk69 expressing cells, in particular in pIIa precursor cells and their progeny. It is thus expected that *ham* would be ectopically expressed in pIIa cells in Ttk69-mutant SOs. Indeed, we observed the presence of three to four Ham-positive cells in 50% of Ttk69-mutant SOs analyzed (SOs inside Ttk69 clones) at 24 h APF, whereas there were no more than two in the control SOs (SOs outside Ttk69 clones) (Figures 7A and B). However, ectopic expression of *ham* could be due to the deregulation of *ham* expression *per se* or to the cousin-cousin cell transformation already described. We assessed *ham* expression in Ttk69-mutant SOs at early stages to determine the mechanism behind its ectopic expression. We observed three Ham-positive cells in Ttk69-mutant SOs composed of four cells as early as 20 h APF, even before the completion of the bristle lineage (Figures 7A and B). These data show that the ectopic expression of *ham* in the absence of Ttk69 is an early event during the cousin-cousin cell transformation. Moreover, we occasionally observed cells positive for both Ham and Pdm1, a specific marker of pIIa descendent cells, in Ttk69-mutant sensory clusters (arrowhead in Figures 7C, C’, n = 4). This signature is consistent with the cells undergoing transformation from outer to inner cells.

**Figure 7.**
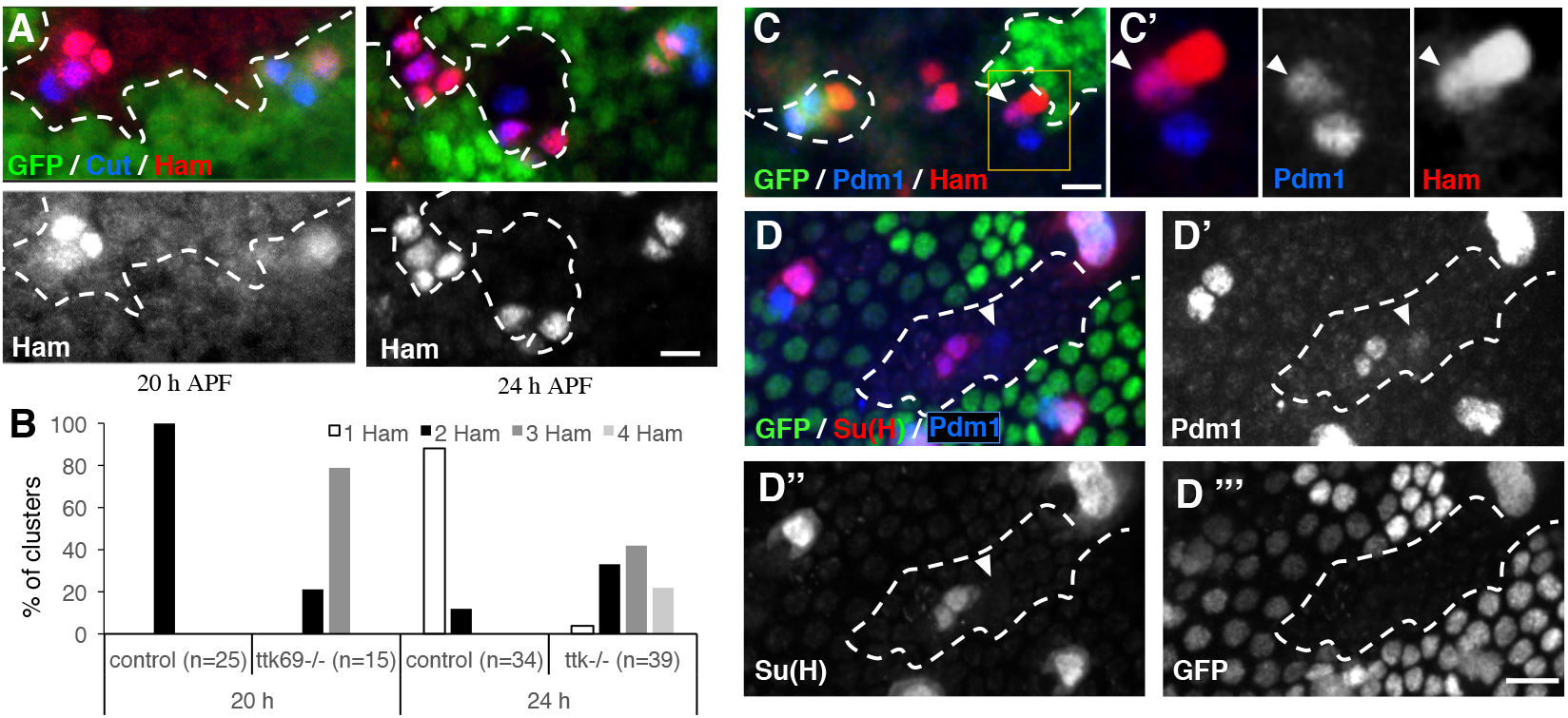
Ttk69 represses *hamlet* expression to maintain the non-neural cell fate. Ttk69 clones outlined by a white line were detected by the absence of GFP (green). (A) *ham* expression (red) in Ttk69-mutant SO at 20 and 24 h APF. Sensory cells in blue. (B) Histogram showing the percentage of SO harboring one (white bars), two (black bars), three (dark grey bars), or four (pale grey bars) Ham-positive cells in control and Ttk69-mutant SO at 20 and 24 h APF. (C) Cell transformation of a Pdm1-positive cell to a Ham-expressing cell. Arrowhead, Ttk69-mutant cell having both Pdm1 (blue in C’) and Ham (red in C’) immunoreactivities. Pupae at 22 h APF. (C’) Higher magnification of the Ttk69-mutant cluster outlined in C. Merged and separate Pdm1 and Ham channels. (D) Recovery of shaft cells. Arrowhead, a Pdm1 positive, Su(H) negative Ttk69-mutant cell in a *ham* heterozygous mutant background. *ham^+/-^* pupae at 28 h APF. D’-D”’, separate Pdm1, Su(H) an GFP channels. Scale bars: 10 μm.

These data suggest that the de-repression of *ham* in pIIa cells in Ttk69-mutants drives cousin-cousin cell transformation, in which pIIa shaft cells adopt a pIIIb cell fate. We tested this possibility by studying whether the reduction of *ham* expression in Ttk69-mutant SOs could restore shaft identity. We performed this analysis late in bristle development, at 28 h APF, when this transformation has already taken place. Under these conditions, we observed Pdm1 positive/Su(H) negative cells, a specific sign of shaft cells, in Ttk69-mutant SOs in a *ham* heterozygous background (4 of 60 Ttk69-mutant SOs analyzed, Figures 7D-D”’, arrowhead). These results show that a reduction in *ham* expression can rescue the formation of shaft cells in the absence of Ttk69. Thus, Ttk69 maintains a non-neural cell fate in pIIa daughter cells by inhibiting the adoption of the inner precursor fate via the repression of *ham* expression.

## Discussion

An important goal in developmental biology is to understand the mechanisms by which cell proliferation and cell-fate acquisition are coupled during organogenesis. Here, we show that Ttk69, a member of the evolutionarily conserved BTB-ZF transcription factors acts as a link between these two processes. We found that Ttk69 is essential for exiting the proliferative progenitor state and conferring a non-neural fate to the progeny during the formation of mechanosensory bristles. Indeed, using mainly clonal analysis, we show that ectopic cell divisions occur independently of changes in cell fate. In addition, we observed that Ttk69-mutant SOs harbor supplementary terminal cells due to cell transformation that generates extra neural progenitor cells. This was associated with upregulation of *cycE*, required for S-phase entry, and the ectopic expression of *hamlet*, a neural determinant. As such, the BTB-ZF transcriptional factor, Ttk69, links cell proliferation and cell fate acquisition in the bristle cell lineage (Figure 8).

**Figure 8.**
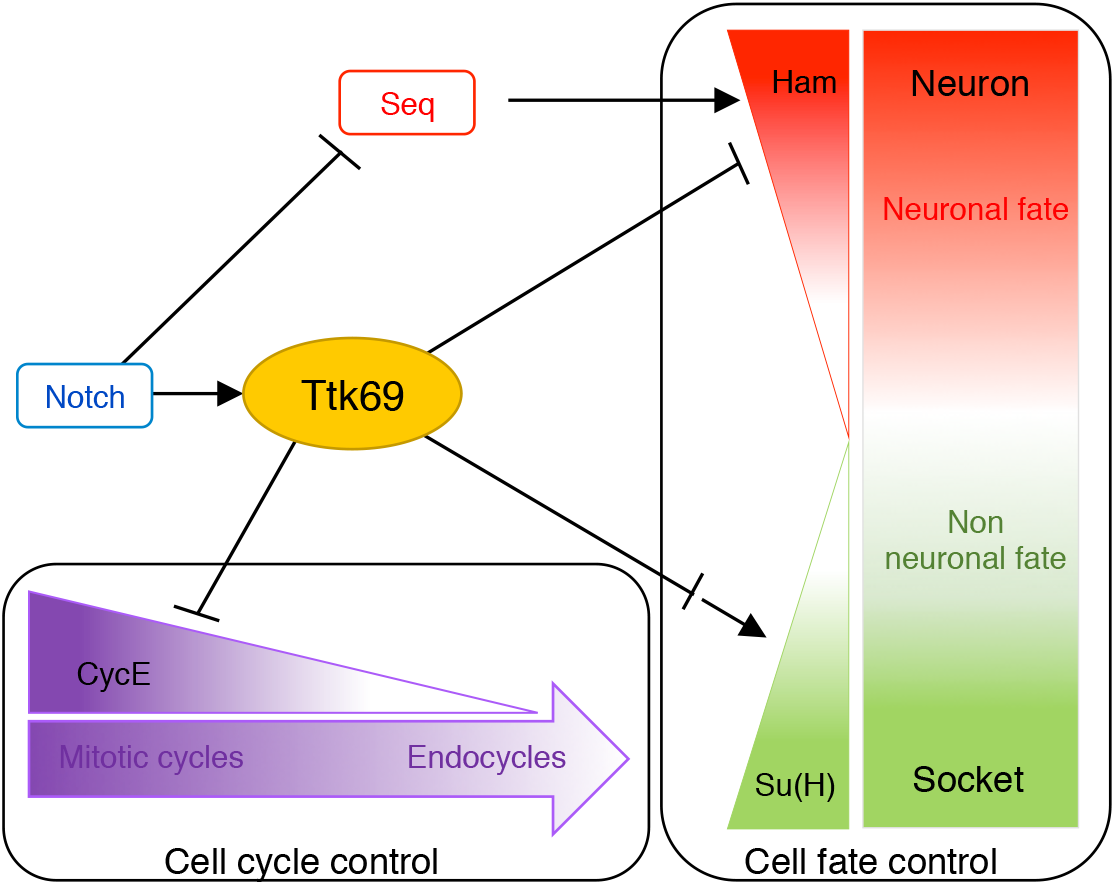
Ttk69 as a central node of a transcriptional regulatory network coordinating terminal cell fate acquisition and the arrest of cell proliferation. In response to N-pathway activation (Guo et al., 1995), Ttk69 (1) downregulates *cycE* expression inducing transition from a mitotic to endocycle mode of the cell cycle and (2) downregulates *ham* and upregulates *Su(H)* expression. Downregulation of *ham* expression prevents the acquisition of neural fate induced by the combined action of Seq and Ham (Andrews et al., 2009; Moore et al., 2004). Upregulation of *Su(H)* expression allows the terminal differentiation of socket cells.

### Ttk69 acts as a dual factor linking cell proliferation and cell fate acquisition

Our results showed that the loss of Ttk69 leads to SOs harboring up to eight terminal cells. Supplementary terminal cells arose from two different mechanisms: an extra division of socket cells and, more importantly, several rounds of extra division due to the re-specification of the presumptive shaft cell into a pIIIb precursor cell. It was previously shown that SOs in complete *ttk* null pupae are composed of only four neurons (Guo et al., 1995). The fact that no ectopic cells were generated when all Ttk isoforms were absent may reflect different kinetics between cell cycle arrest and cell differentiation. *ttk* null cells rapidly acquired an arrested cell cycle and neuronal terminal fate, rather than entering a proliferative precursor state, as after Ttk69 loss of function alone. Thus, the specific effects of Ttk69 on the cell cycle were masked in the *ttk* null mutant. Use of the Ttk69 loss-of-function mutant made it possible to reveal intermediate cell fates, prior to the acquisition of terminal-cell identities. Moreover, the study of mutants that exclusively affect Ttk69 allowed decoupling of the acquisition of cell cycle arrest and cell fate.

The effects of Ttk69 on the core cell cycle machinery were revealed by the ectopic divisions of socket cells observed under Ttk69 loss of function conditions. These cells are already committed to acquire a terminal identity, indicating that such ectopic divisions are not associated with changes in cell fate. This shows that Ttk69 impedes cell cycle progression *per se.* Accordingly, we show that the control of cell-cycle progression by Ttk69 involves transcriptional downregulation of *cycE* expression. Such negative control of *cycE* expression by Ttk69 appears to be a general effect, as it has also been observed in proliferating glia cells (Badenhorst, 2001). Moreover, it has also been shown that Ttk69 represses the expression of *string*, which encodes the phosphatase (Cdc25) essential for G2/M transition in the imaginal eye disc (Baonza et al., 2002). This suggests that Ttk69 represses cell-cycle progression at different phases of the cell cycle. Thus, the induction of ectopic cell divisions in the absence of Ttk69 is probably due to the multiple effects of Ttk69 on cell-cycle progression. The diverse targets of Ttk69 in the cell cycle machinery could explain the involvement of this factor in the transition between different modes of the cell cycle (Jordan et al., 2006; Sun et al., 2008). In SOs, pIIa terminal cells underwent endocycles that require the fine-tuned control of *cycE* expression. Indeed, we have already shown that endocycles are abolished in *cycE* null mutants, occur at very low CycE levels, and become mitotic cycles at high CycE levels (Sallé et al., 2012; Simon et al., 2009). We show here that Ttk69 is involved in a mechanism that limits CycE levels. Thus, Ttk69 is probably involved in the transition from mitotic cell cycles to endocycles in pIIa terminal cells. Similar transitions between two cycling states associated with two different levels of Ttk69 have been observed in ovary epithelial follicular cells during the transition between endocycles to gene amplification (Jordan et al., 2006; Sun et al., 2008). We propose that Ttk69 contributes to the dampening of *cycE* levels, allowing cells to transit throughout different modes of the cell cycle.

In addition to cell proliferation, both pIIa daughter cells were differentially affected under Ttk69-mutant conditions, at the level of terminal differentiation for socket cells and determination for shaft cells. For socket cells, although SOs of adult Ttk69-mutants contain socket cuticular structures, auto-amplification of Su(H) expression is impaired, leading to misshapen sockets (Barolo et al., 2000). For shaft cells, presumptive shaft cells are respecified and acquire a neural progenitor identity due to the mis-expression of *ham*. Ham is normally expressed in pIIIb precursor cells and its overexpression induces the formation of SOs bearing supernumerary cells expressing both ELAV and Pros, such as in pIIIb cells (see Figure 4 in (Moore et al., 2004)). In addition, *ham* loss of function induces the conversion of terminal inner cells into outer cells, suggesting that Ham is essential to acquire the neural precursor fate (Moore et al., 2004). Moreover, we observed that a reduction of *ham* levels in Ttk69-mutant clones decreased shaft re-specification, in agreement with the fact that Ham is an essential regulator of neural precursor fate. Finally, we show that Ttk69 repressed *ham* expression. Overall, these data suggest that re-specification of shaft cells is due to the ectopic expression of *ham* as a consequence of the loss of function of Ttk69. *ham* was also mis-expressed in socket cells, but did not lead to cell transformation. This apparent contradiction may be related with the differential activation of the N-pathway between these two sister cells. Indeed, Ttk69 loss of function induces a cell fate change in shaft cells, a Noff cell. In contrast, Ttk69 loss of function in socket cells, in which the N-pathway is activated as soon as the cells are formed (Remaud et al., 2008), impaired only their late differentiation. These results suggest that early activation of the N-pathway prevents cell fate transformation.

We conclude that Ttk69 is required in terminal cells to represses the neural precursor state by inhibiting both proliferative capacity, by repressing *cycE* expression, and neural fate, by repressing *ham*. This shows that Ttk69 is a central actor in the coordination between cell cycle arrest and cell fate acquisition.

### Tt69 regulates its downstream genes in several ways

In this study, we identified three genes *cycE*, *ham*, and *Su(H)* whose expression was deregulated in Ttk69-mutant clones. *Su(H)* is positively regulated by Ttk69, as revealed by its down-regulation in socket cells in the Ttk69-mutant context. The action of Ttk69 on *Su(H)* enhancer may occur in two different ways. Either Ttk69 acts directly as a transcriptional activator or it represses the expression of an unknown factor, as Ttk69 has always been described as a transcriptional repressor. The time required to express this putative relay factor is consistent with the observation that Ttk69 loss of function affected only late socket cell differentiation. Furthermore, we showed that such Ttk69-mediated regulation occurs through the *Su(H)* 3’end auto-regulatory enhancer, ASE5. It is interesting to note that, the long-lasting high-level of *Su(H)* expression mediated by the auto-regulatory ASE5 loop does not requires N-pathway signaling (Liu and Posakony, 2014). As such, once again Ttk69 affected Notch-independent processes. Further experiments are required to elucidate how Ttk69 activates the auto-amplification of *Su(H)* expression.

In contrast to the downregulation of *Su(H)*, we observed upregulation of *cycE* in the absence of Ttk69, in accordance with the canonical Ttk function as a transcriptional repressor. Surprisingly, Ttk69-mediated *cycE* down-regulation was driven through a promoter domain (fragment D) independently of the canonical AGGAC Ttk69 binding sites (Brown et al., 1991). Moreover, we showed that the Ttk69 zinc-finger domain does not bind subdomains of fragment D (D1, D3), whereas the native Ttk69 protein, present in a late embryonic extract, does. There are two non-exclusive explanations for this observation. Either an uncharacterized Ttk69 domain outside the zinc-finger domain binds directly to the D fragment through a non-canonical binding site or Ttk69 binds indirectly to the *cycE* promoter via an interaction with trans-acting factors. The first explanation is formally possible, but no DNA binding domain has been described in the N-terminal portion of the Ttk69 protein. Nevertheless, it is well known that Ttk69 may bind to other non-canonical binding sites. This is the case for the GTCCTG and TTATCCG sequences in *eve* and *ftz* promoters respectively (Harrison and Travers, 1990; Read and Manley, 1992). However, we observed that Ttk69 continued to downregulate cycE expression in the absence of these non-canonical sites, making this explanation unlikely. In contrast, several lines of evidence support the second explanation. It is known that the activity of Ttk69 can be influenced by the presence of other DNA-binding factors. Thus, the repressive action of Ttk69 depends on interactions with MEP1 and Mi2 proteins, which recruit the ATP-dependent chromatin-remodeling complex (NuRD) (Reddy et al., 2010). Moreover, it has been shown that, although Ttk could bind directly to *eve* promoter repressing *eve* expression (see above), Ttk69 repress *eve* expression independently of their direct binding to DNA, by interacting with GAGA factors through its BTB/POZ domain. When bound to DNA, GAGA zinc-finger factors (Trithorax-like, Trl) activate the transcriptional machinery, but this transcription is inhibited when it is complexed with Ttk69 (Pagans et al., 2004). In bristle sensory cells, RNAi-mediated loss of function of the *MEP1, Mi2*, and *Trl* genes did not affect cell number or cell fates, even when these loss-of-function mutations were analyzed in a sensitized *ttk69* heterozygous background (Figure S5). Despite these results, we favor a model in which another factor is required for Ttk69 to bind the D fragment of the *cycE* promoter. Furthermore, we did not find Ttk binding close to the *cycE* promoter using genome-wide Ttk binding profiles from 0 to 12-h old embryos, published by the modENCODE project. Moreover, *cycE* up-regulation was not observed in genome-wide expression experiments performed in S2 cells treated with dsRNA directed against Ttk69 (Reddy et al., 2010). These data suggest that Ttk69 does not regulate *cycE* expression during early embryonic stages. Thus, Ttk-mediated downregulation of *cycE* expression late in development probably requires cell specific *trans-acting* factors.

The involvement of trans-acting factors would explain the diversity of the Ttk response of particular cells at specific developmental stages. Such diversity mediated by trans-acting factors allows Ttk to regulate the expression of a broad spectrum of genes in bristle sensory cells in response to N-pathway activation, (Figure 8). This is true not only for genes related to cell proliferation, such as *cycE*, but also those controlling cell fate, such as *ham* and *Su(H).* Our data suggest that Ttk69 represses *ham* in pIIa sublineage cells and activates the *Su(H)* autoregulatory loop in socket cells. *cycE* and *ham* regulation occur earlier in this lineage and likely involves the binding of Ttk69 to gene promoters as we have shown for *cycE*. In contrast, Su(H) regulation takes place late in the lineage, implying the probable repression of intermediary relay factors. Trans-acting factors would allow cell-specific responses to Ttk69, whereas intermediary relay factors would allow diversification over time. Thus, since Ttk69 is a N-effector, the mechanism of action of this factor increases the spatial and temporal diversity of the N-pathway cell response.

## Materials and methods

### cycE reporter constructions

To establish transgenic fly strains bearing *cycE* transcriptional reporters, the *eve* promoter of the *eve.p-LacZ.attB* construct (Liberman and Stathopoulos, 2009) was replaced with part of the *cycE* promoter from the *16,4 lacZ* construct engineering by H Richardson (Jones et al., 2000) that contained the full length *cycE* promoter region. 4,6WT fragment covers the Kpn1-Xho1 proximal region of the full-length *cycE* promoter. All other constructs derived from the 4,6WT construct, with deletion of the Kpn1-BssHII and the Ale1-Nco1 fragments in the ΔAC-lacZ construct; deletion of the BssHII-Ale1 fragment in the ΔB-lacZ construct and deletion of the Nco1-Xho1 fragment in the ΔD construct. To generate 4,6m-lacZ promoter bearing mutated Ttk69 binding sites, we used the QuickChange Multi Site-Directed Mutagenesis kit (Stratagene). Experiments were performed following the manufacturer instructions. For each Ttk binding site, specific primer carrying ACTGC sequence (underlined in the sequence) to replace the canonical AGGAC sequence were generated and are listed from the more proximal to the more distal Ttk binding site: 5’GGCATGTTAAA<u>ACTGC</u>TGTTTTAGAACTCAGC3’; 5’GCATATGCATGCC<u>ACTGC</u>AAAGGAGCCG3’; 5’GACGCAGAACA<u>ACTGCA</u>GAAGGCGTCG3’; 5’GATGTCCCAAAAAGTAGAC<u>ACTGC</u>TTTAGCTA3’; 5’TAAATGTTATCAA<u>ACTGC</u>TTGGGGGAGAATTG3’; 5’CGACACATAAGCGC<u>ACTGC</u>TTTATGGG3’ 5’GGGGCACCACTGCATCGAGTATTGAGG3’ 5’CGAGATGCGA<u>ACTGC</u>GATTGCAGCAGC5’. Clones obtained were sequenced and WT fragments of the 4,6WT-lacZ construct were replaced after enzymatic digestion by mutated fragments to generate the 4,6m-lacZ construct. Transgenic flies were generated by *BestGene Inc.* (Chino Hills, CA). All constructs were inserted at the same locus, attP40 on chromosome II, using ΦC31 integrase-based tools, to avoid expression variations due to genomic environment.

### Fly strains

Somatic clones were obtained using the FLP/FRT recombination system (Xu and Rubin, 1993). The *y, w; FRT82B ttk^1e11^/CyO^SM5* line (41754 Bloomington) was crossed with the *y, w, Ubx-FLP; FRT82B ubi-nls::GFP* line (gift of J. Knoblich) to generate *ttk69-null* somatic clones. Somatic *ham* and *seq* clones were generated using the *y, w; FRT40A ham^1^/CyO^SM5* (gift from YN. Jan) crossed with *y, w, Ubx-FLP; FRT40A ubi-nls::GFP and y, w; FRT42D seq^A41^/CyO^SM5 (gift from H. Bellen)* crossed to *y, w; UbxFLP, FRT42A ubi-nls::GFP*, respectively.

Analysis of the Ttk69 loss of function on a *ham* heterozygous background was obtained using a *y, w; FRT40A ham^1^; FRT82B ttk^1e1^/CyO^SM5* crossed with *y, w, UbxFLP; FRT82B ubi-nls::GFP*. Analysis of the Ttk69 loss of function on a *cycE* heterozygous background was obtained using a *y, w; cycE^AR95^; FRT82B ttk^1e11^/CyO^SM5* crossed with *y, w, Ubx-FLP; FRT82B ubi-nls::GFP.* To study Su(H) auto-amplification under Ttk69 loss-of-function conditions, the line y, w; *ASE5-GFP; FRT82B ttk^1e11^/CyO^SM5* was crossed with *y, w, Ubx-FLP; FRT82B ubi-nls::GFP.* To analyze the cycE promoter, the following *cycE* transcriptional reporter lines were used: *(y, w; cycE-4.6-lacZ), (y, w; cycE-4,6m-lacZ), (y, w; cycE-ΔAC-lacZ), (y, w; cycE-ΔB-lacZ) and (y, w; cycE-ΔD-lacZ) and (cycE-D-lacZ corresponding to the 2,9 construct described and gift by H. Richardson).* These cycE transcriptional reporters were analyzed under Ttk69 loss-of-function conditions in pupae at 28 h APF in lines obtained after crossing *y, w, Ubx-FLP; FRT 82B, nls-GFP / TM6 Tb* with the following lines *(4,6WT-lacZ;* FRT82B *ttk^1e11^/* TM6 Tb), *(ΔCm-lacZ;* FRT82B *ttk^1e11^/* TM6 Tb), *(ΔD-lacZ;* FRT82B *ttk^1e11^/* TM6 Tb), *(D-lacZ;* FRT82B *ttk^1e11^/* TM6 Tb)

The GAL4/UAS expression system (Brand and Perrimon, 1993) was used to express the following UAS-constructions in the mechanosensory bristle cell lineage using, as a GAL4 driver, the line *neuralized^p72^-Gal4 (neur^p72^)* (Bellaïche et al., 2001), *UAS-histone H2B::YFP (UAS-H2B::YFP)* (Bellaïche et al., 2001), *UAS-tramtrack69 (UAS-ttk69)* (Badenhorst, 2001), *UAS-hamlet* (UAS-ham), *UAS-sequoia* (UAS-seq) (gift from H. Bellen), *UAS-trl^RNA-valium^* (Bloomington, 40940 and 41582), *UAS-mep1* (Bloomington, 35399; VDRC, 24534) and *UAS-mi-2* (VDRC, 100517). We use the temperature conditional line *UAS-H2B::YFP; neur^p72^, tub-GAL80^ts^* to overexpress these constructs late during SO formation. Fly crosses were carried out at 18°C and pupae were transferred to 30°C at 21 h APF. Pupae were fixed and dissected 7 h later.

Genotypes used in each figures are recapitulated in Supplementary Table S1.

### Immunohistology

Pupal nota were dissected at 17-32 h APF and processed as previously described (Gho et al., 1996). Primary antibodies were: mouse anti-Cut (DSHB, #2B10, 1:500); rabbit anti-β-Galactosidase (Cappel; #55976; 1:500); rabbit anti-GFP (Santa-Cruz Biotechnology, #sc-8334; 1:500); mouse anti-GFP (Roche, N^o^ 11 814 460 001, 1:500), rabbit anti-Pdm1 (gift from T. Préat; École Supérieure de Physique et de Chimie Industrielles, Paris, France; 1:200), rabbit anti-Ttk69 (gift from A. Travers, Medical Research Council, Cambridge, United Kingdom; 1:500), rabbit anti-Ham (gift from YN. Jan, Howard Hughes Medical Institute, San Francisco, USA; 1:500), rabbit anti-Seq (gift from H. Bellen, Baylor College of Medicine, Houston, USA; 1:500); rat anti-ELAV (DSHB, #7E8A10, 1:10); mouse anti-ELAV (DSHB, #9F8A9; 1:100); mouse anti-Pros (gift from C. Doe, Institute of Neuroscience, Eugene, USA; 1:5), rat anti-Su(H) (gift from F Schweisguth, Institut Pasteur, Paris, France 1:500), and rabbit anti-phospho-Histone H3 (Upstate, 06-510, 1:10000). Alexa 488-conjugated secondary antimouse (#A11029), anti-rat (#A11006), anti-rabbit (#A11034), Alexa 568-conjugated secondary anti-mouse (#A11031), anti-rat (#A11077), and anti-rabbit (#A11011) were purchased from Molecular Probes and used at 1:1000. Cy5-conjugated antibodies anti-mouse (#715-175-151), anti-rat (#712-175-153), or anti-rabbit (#711-175-152) were purchased from Jackson Immunoresearch and were used at 1:2000. DNA fragmentation was assayed by TdT-mediated dUTP nick end labelling and performed as previously described (Fichelson and Gho, 2003) (TUNEL kit, Roche Molecular Biochemical). Images were processed with NIH-Image and Photoshop software. All quantifications were done using the ImageJ software (NIH).

### Time lapse microscopy

We performed live imaging of SOs in *neur-H2B::GFP; FRT82B ttk^1e11^* pupae following protocols described previously (Sallé et al., 2012; Simon et al., 2009). The construction *neur-H2B::GFP* (line 22A4, gift of F. Schweisguth) allows the following of SO cells throughout the progression of the bristle lineage. White pupae were collected and aged until 20 h APF at 25°C in a humid chamber before dissection and mounting for imaging. Live imaging data were collected using a spinning disk coupled to an Olympus BX-41 microscope (Roper Scientific, 40X, NA 0.75 objective, CoolSnapHQ2 camera). The temperature of the recording chamber was carefully controlled (±0.1°C) using a homemade Peltier device temperature controller fixed to the microscope stage. Systems were driven by Metamorph software (Universal Imaging). Z-stacks of images were acquired every 3 min and assembled using ImageJ software (NIH). At the end of the movies, pupae were dissected and immunostaining were carried out as described previously. Imaged cells were unambiguously identified by their relative position, nuclear size, and order of birth.

### Tramtrack69 pull down

Experiments were performed either using a batch of *E.* coli-expressed Ttk69::ZF or 20 h old *white drosophila* embryo protein extracts. Ttk69::ZF was expressed from BL21 (DE3) bacteria transformed with a pET15 vector in which the C-terminal fragment (318–641) of Ttk69 protein, containing zinc fingers, was cloned in-frame with a histidine tag (gift from A. Travers). Non-denatured *E. coli* extracts were prepared after a 2 h induction in 0.1mM IPTG and embryo extracts were obtained as described by Wordarz (Wodarz, 2008) (1 mg devitellinized embryos was always extracted in 5 μl lysis buffer to calibrate extraction). DNA templates were generated by PCR using 5’ biotinylated primers. As controls, a *ftz* template was obtained using 5’GGGAGTTGCGCACTTGCTTG and 5’GTGCACGCAACGCTGGTGAG primers, which correspond to the portion of the *fushi-tarazu (ftz) promoter* bearing the canonical AGGAC Ttk69 binding sites and a *RP49* template devoid of this sequence was obtained by PCR using 5’ TGTACTTGGCATCCGCGAG and 5’ CACCAGCACTTCTCCAACAC. Two *cycE* templates were obtained using two set of primers 5’ GCAAGATTATGAATATCTAT – 5’ GTGTGCGCGCATGCGCAACG and 5’ GTTGGATTAACCCTTTCTGG – 5’ AGGATTTAAGTCTCAACTC to cover fragments I and II, respectively, which correspond to the proximal part of the promoter bearing the canonical Ttk69 AGGAC binding site. Im and IIm *cycE* mutated promoters, in which all canonical AGGAC sequences were replaced by a ACTGC sequences, were obtained using the same primers as for fragments I and II of the *cycE* promoter. The three *cycE* templates corresponding to the D fragment are obtained using primers 5’GCTGCCTGCTTGGAGTTGAGAC and 5’GGAAGGTCCAAGACGCATGAC for the D1 fragment, 5’ GTCATGCGTCTTGGACCTTCC and 5’TTATGTGCAGATATTGGGCA for the D2 fragment, and 5’TTATGTGCAGATATTGGGCA and 5’ CTCGAGCTGCCAGCGGCTGC for the D3 fragment. Biotinylated DNA was coupled to streptavidin-coated magnetic beads (M280, Dynal Biotechnology) with 0.1 mg beads per 200 ng DNA, overnight at 4°C. The beads were washed three times in B&W buffer (as recommended by the supplier) and streptavidin-immobilized DNA saturated for 1 h in PBS-20% horse serum before incubation for 1 h with the protein extract in PBS-Triton (0.15%). Protein extracts and beads were separated according to the manufacturer’s instructions and washed four times with 100 mM NaCl/25 mM NaH2PO4. Each fraction was then processed for electrophoreses on SDS polyacrylamide gels. Ttk69::ZF was revealed using mouse anti-Penta-Histidine (1:1000; Qiagen; 34660), whereas Ttk69 protein from embryo extracts was revealed using rabbit anti-Ttk69 (1:4000 gift from A. Travers). Specificity of the Ttk69 antibody was tested by analyzing the immuno-detection of protein extracts from pools of ten 20 h old ttk^1e11^/TM6 and w^1118^ embryos. Anti-β tubulin staining (1:10000, Amersham) was used as a loading control. Revelation was performed using horseradish peroxidase coupled to anti-mouse or anti-rabbit (1:10000, Promega) antibodies coupled to the Super Signal Western blotting detection system (Pierce) according to the manufacturer’s instructions.

## Supporting information

## Acknowledgements

We specially thank YN. Jan, J. Knoblich, H. Bellen, H. Richardson and A. Travers for antibodies. The fly community for fly strains. Heather McLean for critical reading.

## Author contributions

Françoise Simon, Anne Ramat, Jérôme Lacoste, Angélique Burg, Agnès Audibert performed the experiments and carried out data analysis. Agnès Audibert, Sophie Louvet-Vallée, Michel Gho conceptualize, designed experiments and carried out data analysis. Agnès Audibert, Sophie Louvet-Vallée, Michel Gho prepared and edited the manuscript.

## Supplementary data

**Figure S1.**
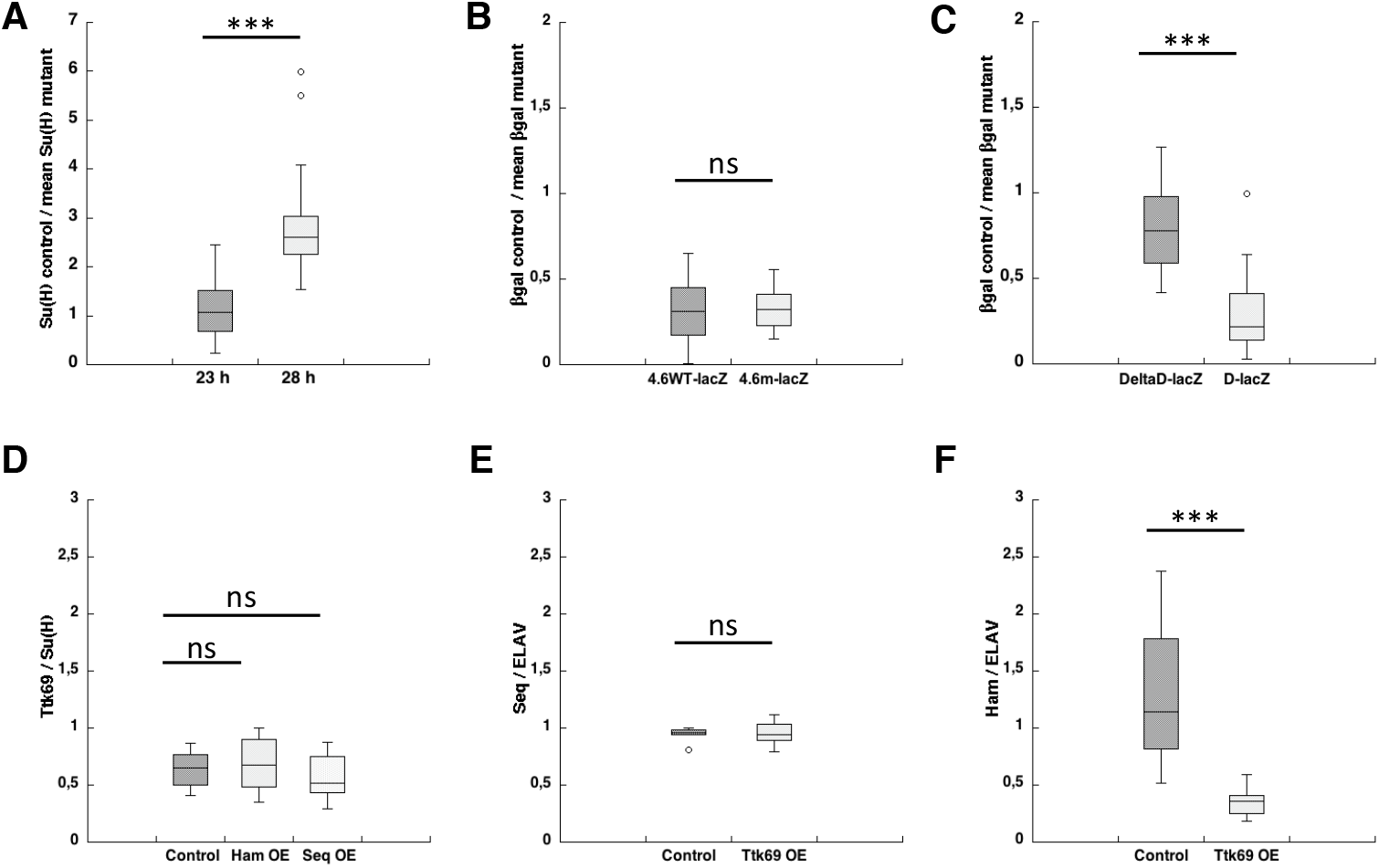
Quantification of immunostaining. Box plots showing quantification of immunostaining under different conditions. (A-C) To compare different experiments, individual quantifications of immunostaining in control conditions were normalized to the mean of similar quantifications in Ttk69-mutant SOs observed in Ttk69 clones. As such, a ratio of 1 indicates that immunostaining in control and Ttk69-mutant SOs is similar, less than one that measurements in control conditions are lower than those observed in Ttk69-mutant cells and conversely when the radio was higher than one. (A) Quantification of data presented in Figures 1F and 1G. Su(H) accumulation in control socket cells at 23h (n = 17 control socket cells and 22 Ttk69-mutant socket cells) and at 28 h APF (n = 38 control socket cells and 33 Ttk69-mutant socket cells). Note that at 23h Su(H) accumulation was similar in both control and Ttk69-mutant socket cells. At 28h APF Su(H) accumulation was 3 times higher in control cells indicating that autoamplification occurred only in the control socket cells. (B) Quantification of data presented in Figures 4C and 4D. β-Gal accumulation in control pIIa cells in the 4.6WT-lacZ fly line (n = 21 control pIIa cells and 30 Ttk69-mutant pIIa cells) and in the 4.6m-lacZ fly line (n = 10 control pIIa cells and 16 Ttk69-mutant pIIa cells). Note that in both cases, ratios are similar and less than 1 indicating that Ttk69 continued to downregulate *cycE* expression when all canonical Ttk69 binding sites were mutated. (C) Quantification of data presented in Figures 5B and 5C. β-Gal accumulation in control pIIa cells in pupae expressing the *ΔD-lacZ* construct (n = 29 control pIIa cells and 30 Ttk69-mutant pIIa cells) and in pupae expressing the *D-lacZ* construct (n = 16 control pIIa cells an d 17 Ttk69-mutant pIIa cells). Note that the *ΔD-lacZ* construct was not downregulated by Ttk69 and behaves similar as to when Ttk69 was absent. Furthermore, the *D-lacZ* construct behaves as control and has a ratio of about 0.3, similar to that observed in *4,6WT-lacZ* and *4,6mlacZ.* (D) Quantification of data presented in Figure S4. Box plot showing quantification of Ttk69 immunostaining. To compare different experiments, individual quantifications of Ttk69 immunostaining in control conditions were normalized to the mean of quantifications of Su(H) accumulation observed in socket cells in control (n = 10) and after Ham (n= 12) or Seq (n= 13) overexpression. Note that ratios were unchanged between this three conditions showing that the overexpression of Seq or Ham did not affect Ttk69 expression. (E, F) Quantification of data presented in Figure 6. Box plots showing quantification of Seq immunostaining (D, n = 5) and Ham immunostaining (E, n = 11). To compare different experiments, individual quantifications of immunostaining in control conditions were normalized to the mean of similar quantifications of ELAV staining observed in neurons in control and after Ttk69 overexpression. Note that Ham but not Seq accumulation was reduced after Ttk69 overexpression reflecting a specific downregulation of *ham* expression by Ttk69. ***: p < 0,001, ns: not significant (Mann-Whitney test).

**Figure S2.**
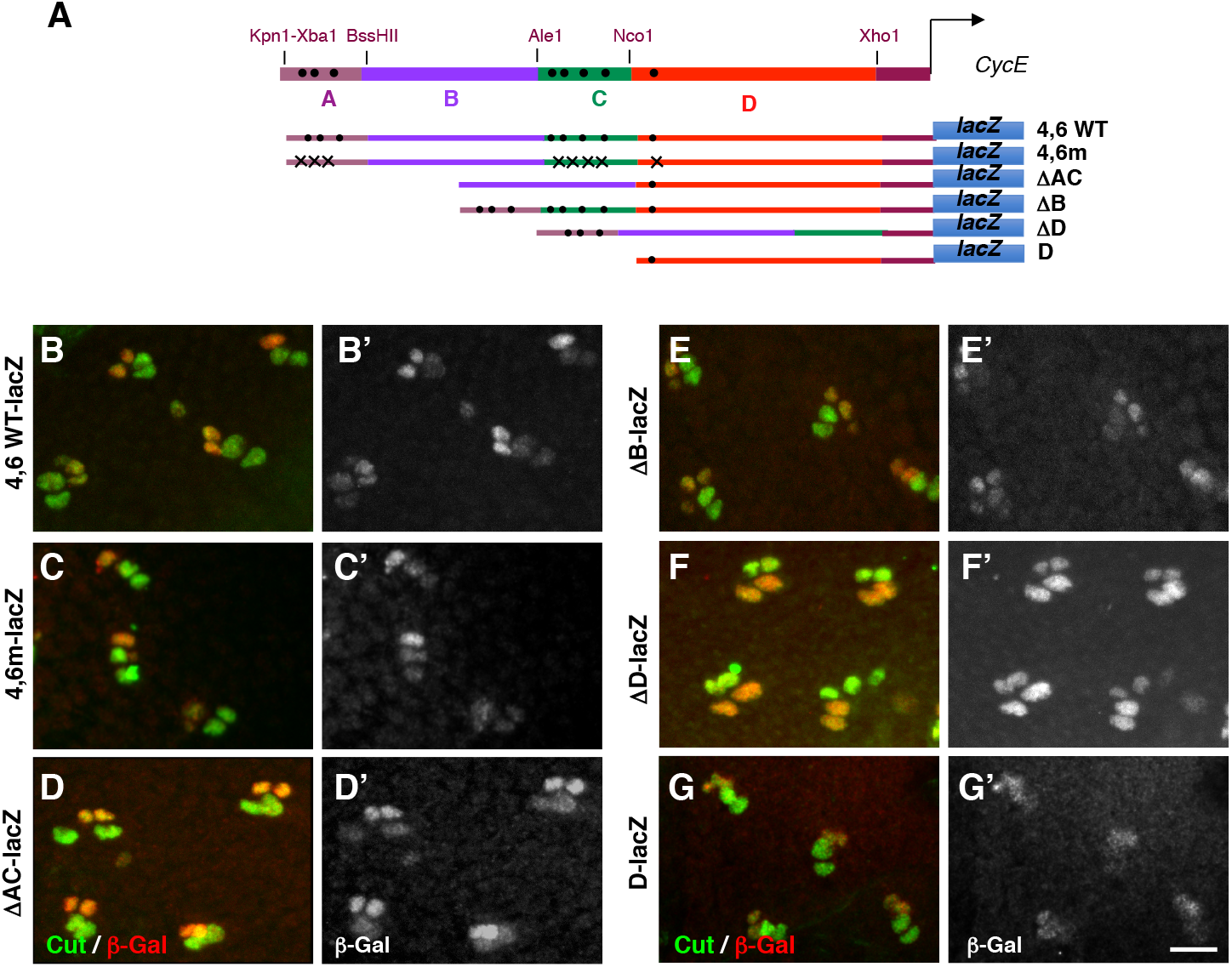
Expression pattern of *cycE* transcriptional reporters in SOs. (A) Diagram of *cycE* transcriptional reporters aligned to the *cycE* promoter at the top. The localization of four specific regions (A to D) is depicted. Black dots indicate localization of canonical AGGAC Ttk-binding sites. 4.6WT *cycE* transcriptional reporter bearing A to D regions; 4.6m, bearing A to D regions in which the eight AGGAC binding sites are mutated; ΔAC, bearing the B and D regions; ΔB, bearing the A, C, and D regions; ΔD, bearing the A to C regions and D bearing only the D region. (B-G’) Expression pattern of the six transcriptional reporters described in (A). Pupae at 28 h APF. Sensory cells were identified by Cut immunoreactivity (green) and expression of *cycE* transcriptional reporters was revealed by β Gal immunoreactivity (B’ to G’ and in red in B to G). Note that the *ΔD-lacZ* construct is expressed in all sensory cells, whereas the expression of the other constructs is higher in the inner cells than in the outer cells. Scale bars: 10 μm.

**Figure S3.**
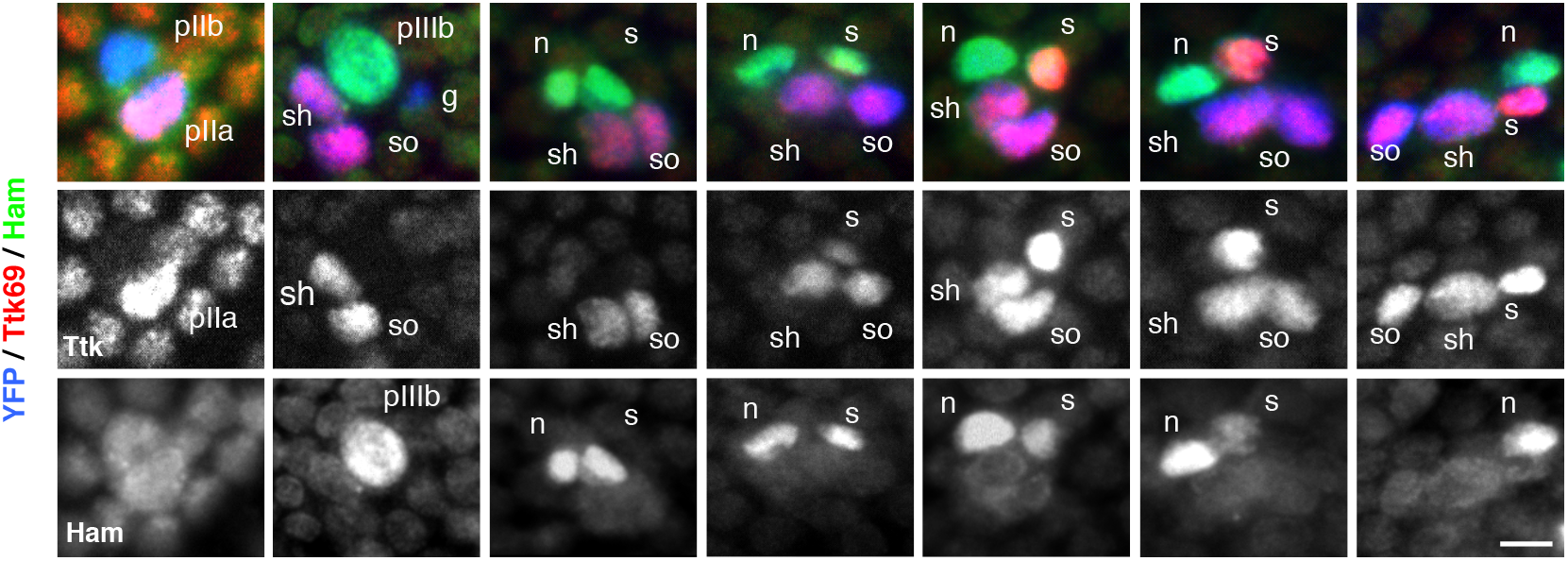
Complementary expression pattern of Ttk69 and Ham proteins in bristle sensory cells. Ttk69 and Ham expression in bristle sensory cells at progressive stages of development from 18 to 28 h APF. Sensory cells are shown in blue (YFP staining), Ttk69-positive cells in red (shown as a separate channel in the middle panels), and Ham-positive cells in green (shown as a separate channel in the bottom panels). The sensory cells shown are the precursor cells: pIIb, pIIa, and pIIIb and the terminal cells: glial cells (g), sheath cells (s), neurons (n), shaft cells (sh), and socket cells (so). Ttk69 is first detected in pIIa cells and their daughter cells and later in the sheath cells. Ham is first detected in pIIIb cells and their daughter cells, before disappearing from the sheath cell when Ttk69 appears in this cell. Scale bars: 5 μm.

**Figure S4.**
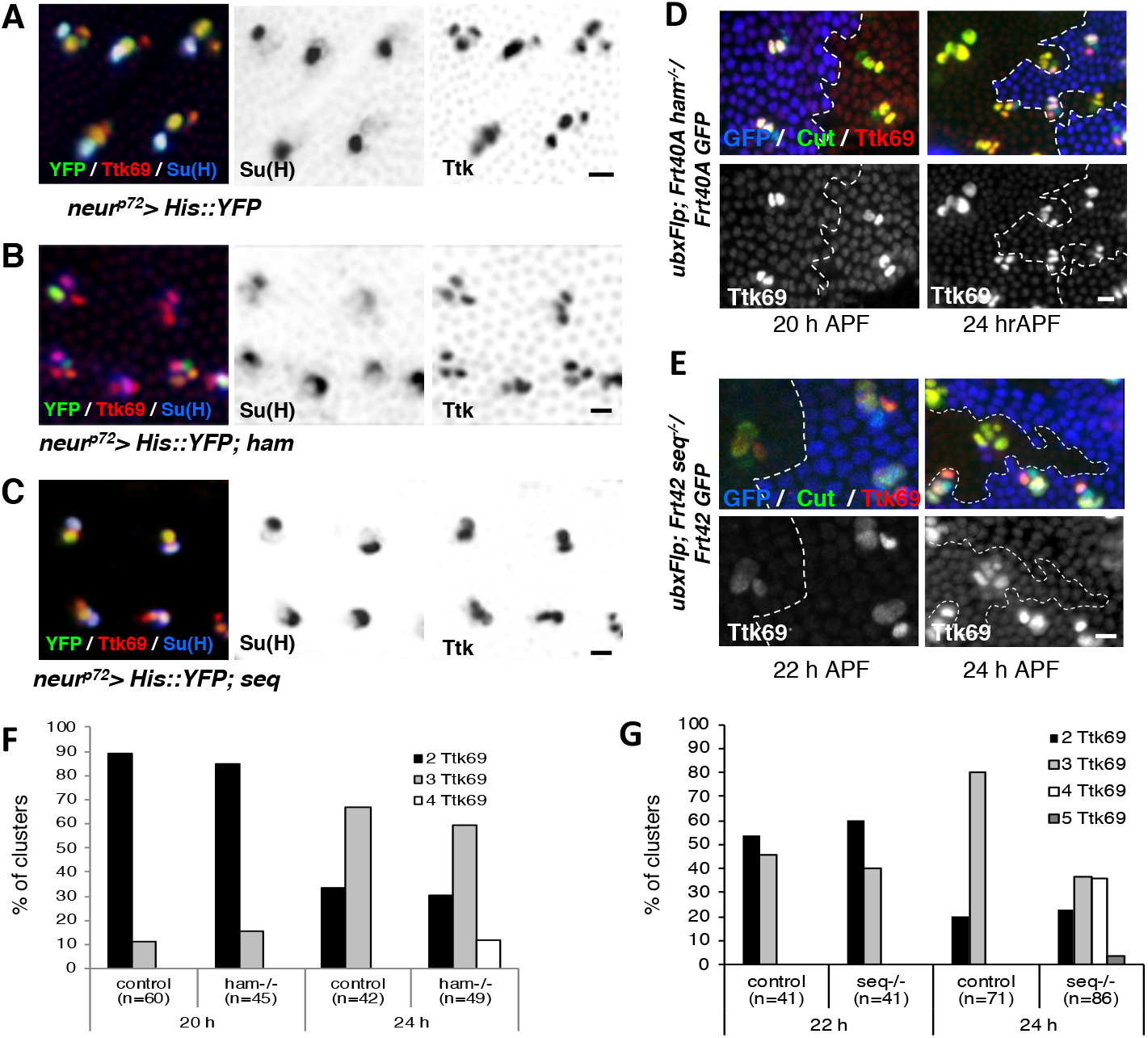
Hamlet and Sequoia do not regulate *ttk69* expression. (A-C) Ttk69 expression under *ham* and *seq* over-expression conditions. Pupae were maintained at 18°C from 0 to 19h APF, shifted to 30°, and analyzed 7 h later. Sensory cells were visualized by YFP expression. Socket cells were identified by Su(H) (blue) and Ttk69 (red) immunoreactivity. Individual, Su(H) and Ttk69 channels are shown in inverted fluorescence (Middle and right panels). Note that, under these conditions, no cell fate transformation occurred, assessed by the normal expression of Su(H) in socket cells. (D, E). Ttk69 expression in clones null for *ham* and *seq* at two developmental times. Clones null for *ham* (D) and *seq* (E) were detected by the absence of GFP (blue, white dashed lines indicating the border of the clones) in pupae at 20, 22, and 24 h APF. Sensory cells were identified by Cut (green) and Ttk69 (red) immunoreactivity (shown also in separate panels). Note that no modification of Ttk expression is observed in mutant SOs (SOs inside the clone) relative to control SOs (SOs outside the clone) at earlier developmental times. Modifications observed at later timepoints are related to *ham* or *seq*-mediated changes in cell identity, rather than the direct action of *ham* or *seq* on *ttk* expression. (F, G) Quantification of experiments shown in D and E. Histograms showing the percentage of SOs harboring two to five Ttk-positive cells in SOs in control (SOs outside clones) and mutant (inside clones) conditions for *ham* (F) and *seq* (G) clones at 20, 22, and 24 h APF. The percentage of clusters with two to five (grey bars) Ttk69 cells is indicated. Note that no significant modifications are observed at earlier developmental times. Scale bars: 10 μm.

**Figure S5.**
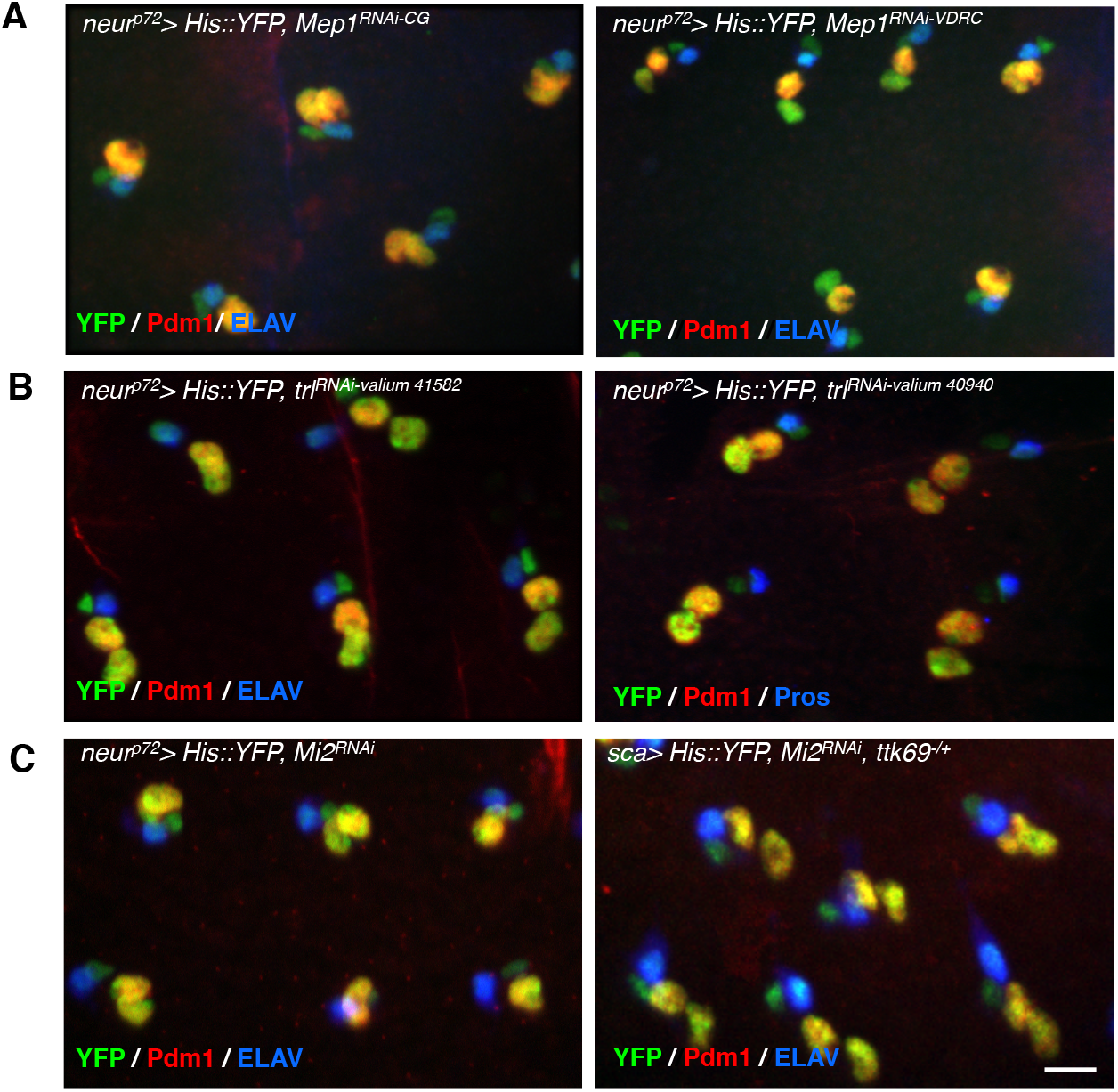
Known partners of Ttk69 do not affect bristle-cell lineage. Identity of SO cells under conditions of RNAi-mediated loss of function for *Mep1* (A), *trl* (GAGA factor) (B), and for *Mi2* (C) in wild-type (left) and Ttk69 heterozygous mutant backgrounds (right). Sensory cells are shown in green (GFP staining). Socket and shaft cells were detected by their specific accumulation of Pdm1 (red), and neurons or sheath cells by ELAV or Pros immunoreactivity (blue). Note that, in all contexts, SOs contained four terminal cells with two outer cells and two inner cells as in control organs. Scale bars: 10 μm.

**Table S1.**
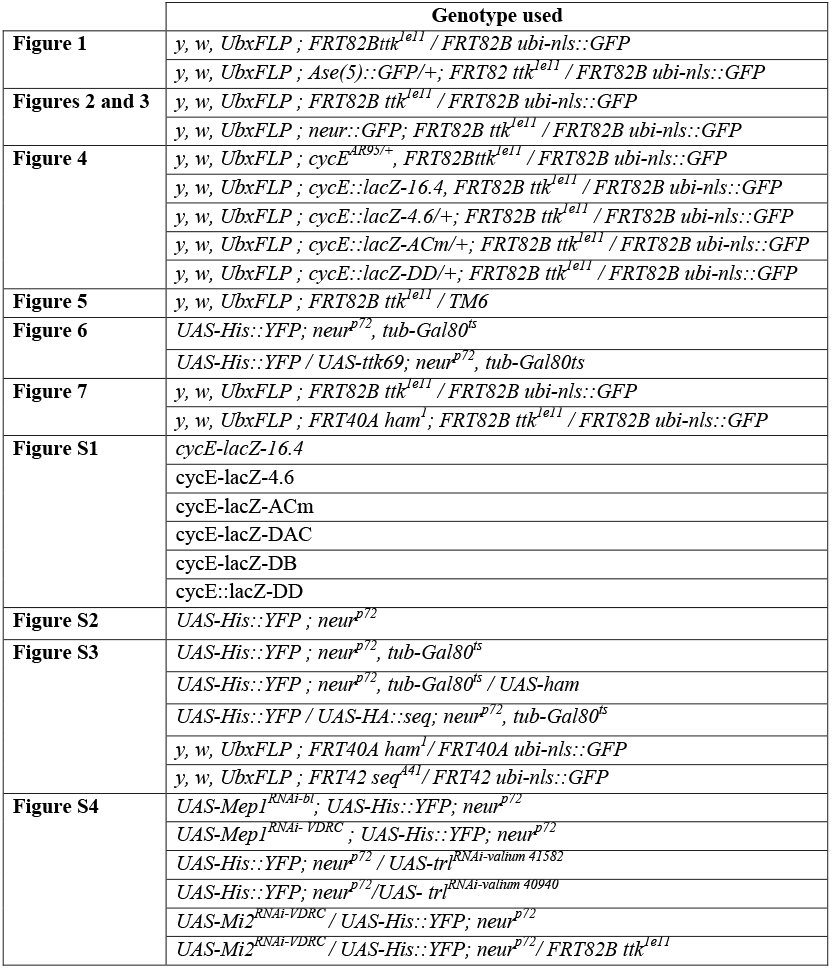
Fly genotypes. Fly genotypes used in each figure.

