## Supplementary material for "The BTB-ZF transcription factor Tramtrack69 shapes neural cell lineages by coordinating cell proliferation and cell fate"

| | **Genotype used** |
|---|---|
| **Figure 1** | y, w, UbxFLP ; FRT82Bttk$^{1e11}$ / FRT82B ubi-nls::GFP |
| | y, w, UbxFLP ; Ase(5)::GFP/+; FRT82 ttk$^{1e11}$ / FRT82B ubi-nls::GFP |
| **Figures 2 and 3** | y, w, UbxFLP ; FRT82B ttk$^{1e11}$ / FRT82B ubi-nls::GFP |
| | y, w, UbxFLP ; neur::GFP; FRT82B ttk$^{1e11}$ / FRT82B ubi-nls::GFP |
| **Figure 4** | y, w, UbxFLP ; cycE$^{AR95/+}$, FRT82Bttk$^{1e11}$ / FRT82B ubi-nls::GFP |
| | y, w, UbxFLP ; cycE::lacZ-16.4, FRT82B ttk$^{1e11}$ / FRT82B ubi-nls::GFP |
| | y, w, UbxFLP ; cycE::lacZ-4.6/+; FRT82B ttk$^{1e11}$ / FRT82B ubi-nls::GFP |
| | y, w, UbxFLP ; cycE::lacZ-ACm/+; FRT82B ttk$^{1e11}$ / FRT82B ubi-nls::GFP |
| | y, w, UbxFLP ; cycE::lacZ-DD/+; FRT82B ttk$^{1e11}$ / FRT82B ubi-nls::GFP |
| **Figure 5** | y, w, UbxFLP ; FRT82B ttk$^{1e11}$ / TM6 |
| **Figure 6** | UAS-His::YFP; neur$^{p72}$, tub-Gal80$^{ts}$ |
| | UAS-His::YFP / UAS-ttk69; neur$^{p72}$, tub-Gal80ts |
| **Figure 7** | y, w, UbxFLP ; FRT82B ttk$^{1e11}$ / FRT82B ubi-nls::GFP |
| | y, w, UbxFLP ; FRT40A ham$^{1}$; FRT82B ttk$^{1e11}$ / FRT82B ubi-nls::GFP |
| **Figure S1** | cycE-lacZ-16.4 |
| | cycE-lacZ-4.6 |
| | cycE-lacZ-ACm |
| | cycE-lacZ-DAC |
| | cycE-lacZ-DB |
| | cycE::lacZ-DD |
| **Figure S2** | UAS-His::YFP ; neur$^{p72}$ |
| **Figure S3** | UAS-His::YFP ; neur$^{p72}$, tub-Gal80$^{ts}$ |
| | UAS-His::YFP ; neur$^{p72}$, tub-Gal80$^{ts}$ / UAS-ham |
| | UAS-His::YFP / UAS-HA::seq; neur$^{p72}$, tub-Gal80$^{ts}$ |
| | y, w, UbxFLP ; FRT40A ham$^{1}$/ FRT40A ubi-nls::GFP |
| | y, w, UbxFLP ; FRT42 seq$^{A41}$/ FRT42 ubi-nls::GFP |
| **Figure S4** | UAS-Mep1$^{RNAi\text{-}bl}$; UAS-His::YFP; neur$^{p72}$ |
| | UAS-Mep1$^{RNAi\text{- }VDRC}$ ; UAS-His::YFP; neur$^{p72}$ |
| | UAS-His::YFP; neur$^{p72}$ / UAS-trl$^{RNAi\text{-}valium\ 41582}$ |
| | UAS-His::YFP; neur$^{p72}$/UAS- trl$^{RNAi\text{-}valium\ 40940}$ |
| | UAS-Mi2$^{RNAi\text{-}VDRC}$ / UAS-His::YFP; neur$^{p72}$ |
| | UAS-Mi2$^{RNAi\text{-}VDRC}$ / UAS-His::YFP; neur$^{p72}$/ FRT82B ttk$^{1e11}$ |

**Table S1. Fly genotypes.** Fly genotypes used in each figure.
